# Concerted 2-5A-Mediated mRNA Decay and Transcription Reprogram Protein Synthesis in dsRNA Response

**DOI:** 10.1101/484675

**Authors:** Sneha Rath, Eliza Prangley, Jesse Donovan, Kaitlin Demarest, Ned S. Wingreen, Yigal Meir, Alexei Korennykh

## Abstract

RNA degradation by RNase L during 2-5A-mediated decay (2-5AMD) is a conserved mammalian stress response to viral and endogenous double-stranded RNA (dsRNA). 2-5AMD onsets rapidly and facilitates a switch of protein synthesis from homeostasis to production of interferons (IFNs). To understand the mechanism of this protein synthesis reprogramming, we examined 2-5AMD in human cells. 2-5AMD triggers polysome collapse characteristic of a translation initiation defect, but translation initiation complexes and ribosomes purified from the translation-arrested cells remain functional. Using spike-in RNA-seq we found that basal messenger RNAs (mRNAs) rapidly decay, while mRNAs encoding IFNs and IFN-stimulated genes evade 2-5AMD and accumulate. The IFN evasion results from the combined effect of better mRNA stability and positive feedback amplification in the IFN response. Therefore, 2-5AMD and transcription act in concert to revamp the cellular mRNA composition. The resulting preferential accumulation of innate immune mRNAs establishes “prioritized” synthesis of defense proteins.

## Introduction

The innate immune system is activated without the delay needed for antibody production and provides an immediate defense barrier against infections and out-of-control host cells posing danger. In higher vertebrates, the innate immune system relies on interferon (IFN) signaling coupled with a vertebrae-specific pathway of regulated RNA degradation, 2-5A-mediated decay (2-5AMD) (Chakrabarti et al., 2011; Cooper et al., 2014b; Donovan et al., 2017; Rath et al., 2015). 2-5AMD is activated in the presence of double-stranded RNA (dsRNA), a pathogen-associated and damage-associated molecular pattern that signals the presence of viruses (Li et al., 2016) and pathologic derepression of endogenous repeat elements of the host (Chiappinelli et al., 2015; Leonova et al., 2013; Li et al., 2017).

2-5AMD requires the coordinated activity of 2-5A oligonucleotide synthetases (OASs) and the pseudokinase-endoribonuclease, RNase L. The action of the OASs closely parallels that action of a structurally similar dsDNA sensor cGAS. cGAS synthesizes a second messenger cGAMP (cyclic-G_2′,5′_A_3′,5′_p) to activate the IFN response via the cGAMP receptor, STING (Civril et al., 2013). The OASs use the same mechanism of regulation, but function as sensors of dsRNA (Civril et al., 2013; Donovan et al., 2013). The OASs synthesize the second messenger 2-5A (5′-ppp-A_2′p5′_A(_2′p5′_A)_n≥0_) to activate RNA decay by the 2-5A receptor, RNase L (Chakrabarti et al., 2011; Donovan et al., 2013; Donovan et al., 2015).

RNase L is a mammalian pseudokinase-endoribonuclease uniquely related to the kinase-RNase Ire1 in the unfolded protein response (Korennykh et al., 2009; Lee et al., 2008; Zhou et al., 2000). 2-5A binding to the ankyrin-repeat sensor domain of RNase L facilitates its dimerization and high-order oligomerization, which provides the switch for activation of RNA cleavage by 2-5AMD (Han et al., 2014; Han et al., 2012). Upon activation, RNase L cleaves single-stranded RNA molecules at UN^N sites (Floyd-Smith et al., 1981; Han et al., 2014). The prevalence of this short motif results in 2-5AMD sensitivity of many tRNAs, rRNAs, mRNAs, Y-RNAs and vault RNAs (Cooper et al., 2014b; Donovan et al., 2017). During homeostasis, 2-5AMD regulates adhesion and migration activity of mammalian cells (Banerjee et al., 2015; Rath et al., 2015). Upon acute activation by dsRNA, 2-5AMD arrests global translation by an immediate and poorly understood mechanism (Alisha Chitrakar et al., 2018; Donovan et al., 2017). In mammalian cells, dsRNA inhibits translation not only via 2-5AMD, but also via phosphorylation of the translation initiation factor eIF2α by the serine-threonine kinase protein kinase R (PKR) (Fig. 1A). 2-5AMD and the PKR pathway are separated in time and in A549 human lung epithelial cells, 2-5AMD is the only mechanism responsible for the rapid translational inhibition (Donovan et al., 2017).

**Figure 1.**
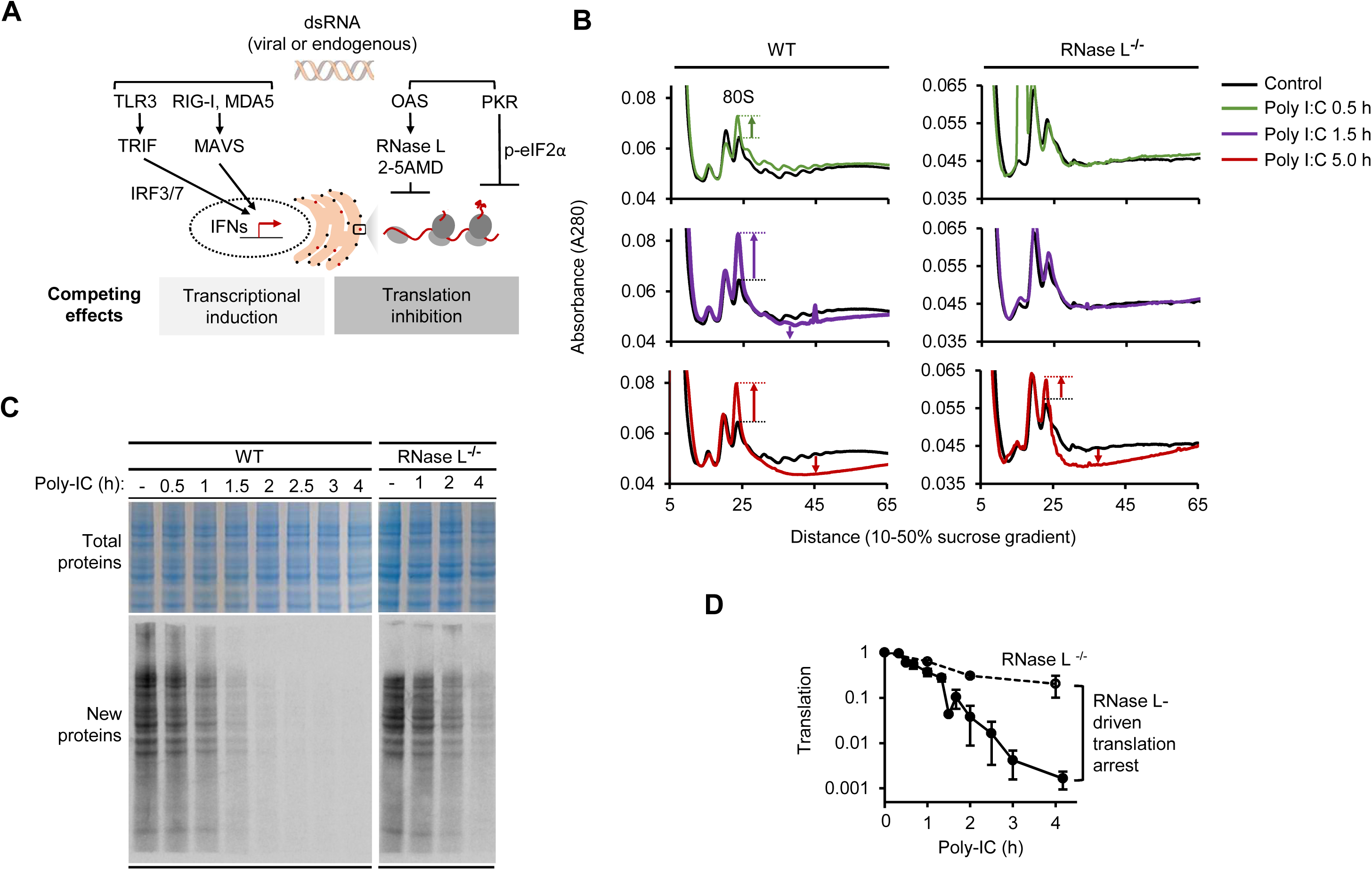
The effect of 2-5AMD and RNase L activation on polysomes in A549 human cells. (A) Schematic overview of the dsRNA sensing pathways and the location of 2-5AMD. (B) Polysome sedimentation profiles in WT and RNase L^-/-^ cells during poly I:C treatment. (C) Translation activity in WT and RNase L^-/-^ cells after poly I:C transfection measured using ^35^S metabolic labeling. Total proteins stained with coomassie show lane loading. Independent measurements using an orthogonal readout are shown in related Fig. S1. (D) Quantification of new protein synthesis, normalized to loading control. Error bars are S.E. from three biological replicates.

Initial studies of 2-5AMD in rabbit reticulocyte lysate (RRL) reported polysome disaggregation and degradation of mRNAs (Clemens and Williams, 1978). RRL does not have transcription, which limits biologic interpretations of these results: in the cell-free system, the loss of mRNAs and polysomes due to a non-specific RNase is inevitable. In live cells, 2-5AMD has sufficient potency to cause degradation of 18S and 28S rRNAs. The characteristic rRNA cleavage pattern provides a reliable readout of RNase L activation (Donovan et al., 2017; Malathi et al., 2007). Furthermore, 2-5AMD causes degradation of tRNA-His and tRNA-Pro, as well as multiple mRNAs in live cells (Al-Ahmadi et al., 2009; Donovan et al., 2017; Le Roy et al., 2007; Rath et al., 2015). Due to the complexity of the RNA degradation program, the arrest of global translation by 2-5AMD could not be linked to any specific RNA (Alisha Chitrakar et al., 2018; Donovan et al., 2017).

Shortly after activation of 2-5AMD, dsRNA triggers the transcriptional IFN response, leading to upregulation of innate immune mRNAs (Fig. 1A) (Alisha Chitrakar et al., 2018; Kawai et al., 2005; Liu et al., 2008). While global translation remains silenced by actively ongoing 2-5AMD, the mRNAs encoding the interferons IFN-β (type I) and IFN-λ (type III) bypass the global inhibition and are actively translated (Alisha Chitrakar et al., 2018). Here we employ cell biology, proteomics, transcriptomics and modeling approaches to determine the translation reprogramming mechanism of 2-5AMD and to explain how innate immune mRNAs can evade the translational arrest.

## Results

### 2-5AMD Inhibits Translation Initiation without Disrupting Cap-binding Complex and 40S Subunit Loading

To begin deciphering the mechanism of protein synthesis regulation 2-5AMD, we examined whether it affects polysomes in WT and RNase L^-/-^ cells by sucrose gradient sedimentation. In the presence of immunogenic dsRNA (poly-I:C), the 80S monosome peak became dominant within 30 minutes, whereas polysomes progressively disassembled (Fig. 1B). In RNase L^-/-^ cells that experience weak translation inhibition by PKR, the polysome profiles did not change until five hours, and the 80S peak never dominated. The changes in the polysome profiles were accompanied by a global, RNase L-dependent arrest of translation (Fig. 1C-D; Fig. S1A-B). The loss of polysomes and accumulation of the 80S monosomes during 2-5AMD is a signature of inhibited initiation of capped mRNAs. A similar polysome change takes place upon deletion of the RNA helicase DHX29 involved in recognition of the 5′-untranslated region (5′-UTR) (Parsyan et al., 2009) or inhibition of the mammalian target of rapamycin (mTOR) that maintains translation initiation active (Gandin et al., 2014). The 80S species that form upon DHX29 and mTOR defects are non-translating, as are the 80S species formed during 2-5AMD (Fig. 1C; Fig. S1A). Non-translating 80S species devoid of mRNA form readily in A549 cells following translation release with puromycin (Fig. S1C). Puromycin-induced and 2-5AMD-induced 80S monosomes are stable only at ∼100 mM KCl, but dissociate at 500 mM KCl (Fig. S1C-D), as expected for vacant ribosomes (van den Elzen et al., 2014).

Inhibition of mTOR kinase is a common strategy to arrest bulk translation during stress responses (Hsieh et al., 2012; Zoncu et al., 2011), suggesting that inhibition of translation initiation by 2-5AMD could depend on mTOR. To test this link, we assessed mTOR activity by measuring phosphorylation of the translation initiation factor 4E-BP1 whose phosphorylation by mTOR is required for translation initiation (Feldman et al., 2009). Activation of 2-5AMD did not affect 4E-BP1 phosphorylation, whereas a control treatment with the small molecule mTOR inhibitor INK128 (Feldman et al., 2009) worked (Fig. 2B). Considering that 2-5A and INK128 inhibited bulk translation comparably but with different effects on 4E-BP1 phosphorylation (Fig. 2A-B), our data suggest that 2-5AMD inhibits translation initiation independently from mTOR.

**Figure 2.**
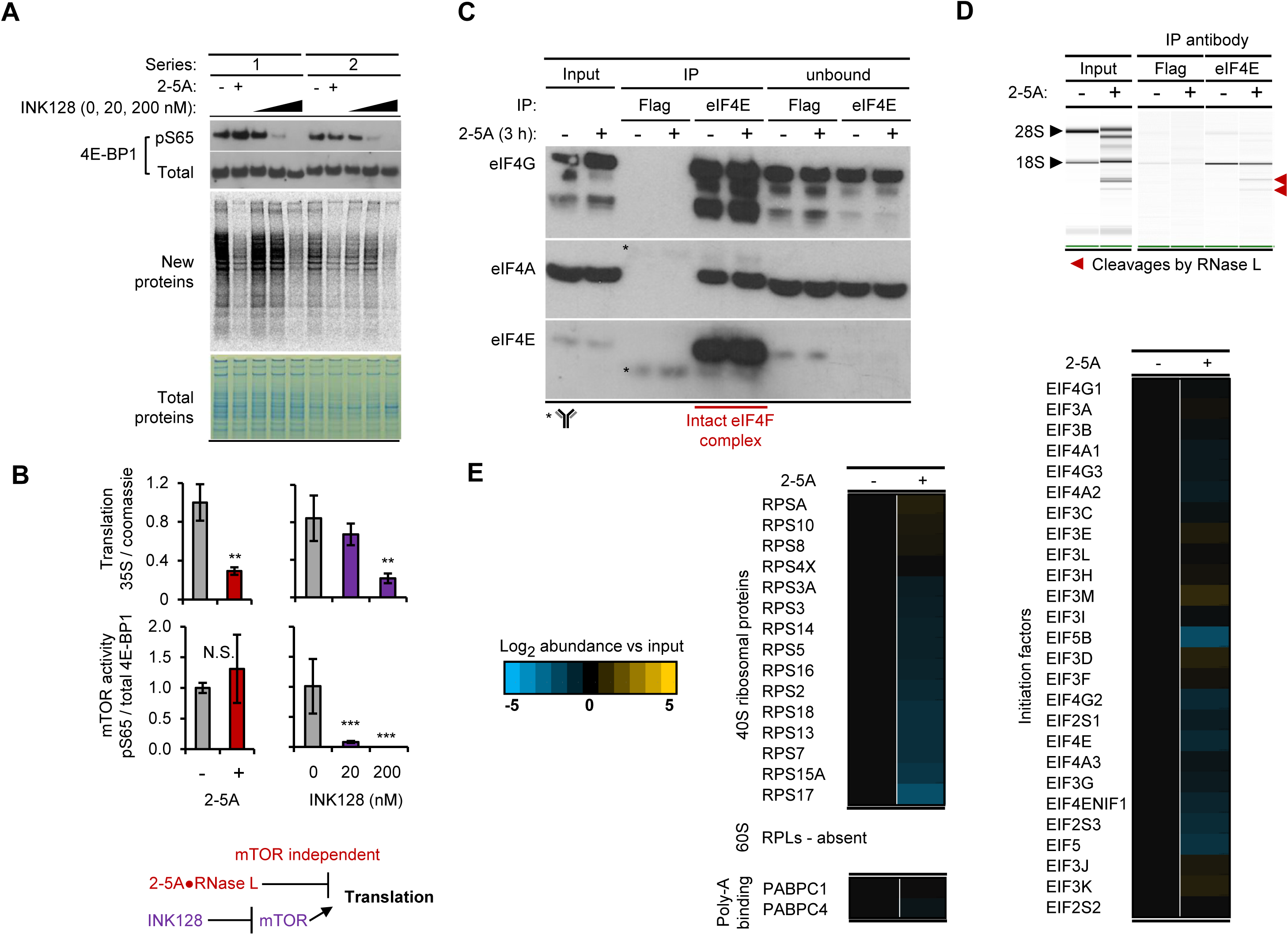
Translation initiation status during 2-5AMD. (A) Activity of mTOR kinase monitored by western blot for phosphorylated 4E-BP1. Matching translational activity was measured by ^35^S metabolic labeling. The small molecule mTOR inhibitor INK128 is used as a control. (B) Quantification of the gels in (A). (C) Western blot analysis of core components of the cap-binding eIF4F complex: eIF4E, eIF4A and eIF4G. Data from control (Flag) and eIF4E immunoprecipitation (IP) experiments are shown with and without 2-5A treatment. (D) BioAnalyzer NanoChip profiling of rRNA in the samples in (C). (E) Mass spectrometry analysis of proteins that co-immunoprecipitate with the cap-binding translation initiation factor eIF4E (Table S1).

To test whether 2-5AMD disrupts assembly of cap binding initiation complexes, we pulled down the cap-binding initiation factor eIF4E and examined its association with the key partner factors eIF4A and eIF4G that together form the eIF4F complex. This tripartite complex was readily identified using the pulldown and remained unchanged by 2-5AMD (Fig. 2C). Total RNA profiling by NanoChip revealed that eIF4E additionally pulled down the 40S ribosomal subunit both in naïve cells and in cells with activated 2-5AMD, suggesting normal loading of the small subunit. As expected, 18S and 28S rRNAs were degraded during 2-5AMD and exhibited the characteristic pattern of RNase L activity (Fig. 2D). The 18S rRNA from the 40S subunit pulled down with eIF4E following 2-5AMD was cleaved as in the input rRNA. Thus, binding of the core components of the translation initiation complex is not disrupted during 2-5AMD.

To examine the initiation complex more completely, we performed mass spectrometry (MS) of proteins that co-purify with eIF4E (Fig. S2). This analysis detected ribosomal proteins from the small subunit, translation initiation factors and poly-A binding proteins, but not proteins from the large ribosomal subunit, as expected due to polysome run off. The identified components did not change in response to 2-5A (Fig. 2E; Table S1), indicating 2-5AMD does not block the cap-binding complex and cap-mediated loading of the small ribosomal subunit.

### 2-5AMD Degrades 18S and 28S rRNAs but Leaves Ribosomes Functional

2-5AMD does not interfere with the initiation step. To further define the mechanism, we analyzed the hallmark substrate of RNase L, the ribosome, and examined whether 2-5AMD inhibits global translation by directly affecting the ribosomal translation activity. 18S and 28S rRNAs are both cleaved by RNase L (Fig. 2D) (Donovan et al., 2017; Malathi et al., 2007). Although a high-resolution structure of the human 80S ribosome has become available (Khatter et al., 2015), mechanistic understanding of how rRNA cleavage by 2-5AMD could affect the ribosome is critically limited by the absence of reliable mapping of the cleavage sites. Based on primer extension analysis it has been proposed that RNase L cleaves nucleotides 4032, and to a smaller extent 4031, in 28S rRNA (Iordanov et al., 2000). In contrast, subsequent RNA-seq analyses of cleaved rRNAs captured using *Arabidopsis* tRNA ligase did not detect cleavage at either of the 28S rRNA sites, but observed predominantly RNase L-independent 18S/28S rRNA background cleavage events (Cooper et al., 2014b).

To *de novo* identify the cleavage sites in human rRNA, we adapted RtcB RNA-seq suitable for single-nucleotide resolution mapping (Donovan et al., 2017). RtcB RNA-seq analysis of rRNA from cells with activated 2-5AMD (Fig. S3A) revealed cleavage sites with a UN^N consensus (Fig. 3A) that distinguishes the sequence-specific activity of RNase L (Table S2) (Donovan et al., 2017; Han et al., 2014). We analyzed the data to identify predominant sites that simultaneously exhibit induction and high read count in the 2-5AMD sample, as shown in Figure 3A. A single high-scoring site 771 was detected in 18S rRNA and two dominant sites, 1056 and 4032, were detected in 28S rRNA. Our analysis detected both 28S rRNA sites 4031 and 4032 identified previously by primer extension assay (Iordanov et al., 2000), providing important validation for the RtcB RNA-seq approach. The dominant sites 771 (18S) and 1056 (28S) were detected for the first time.

**Figure 3.**
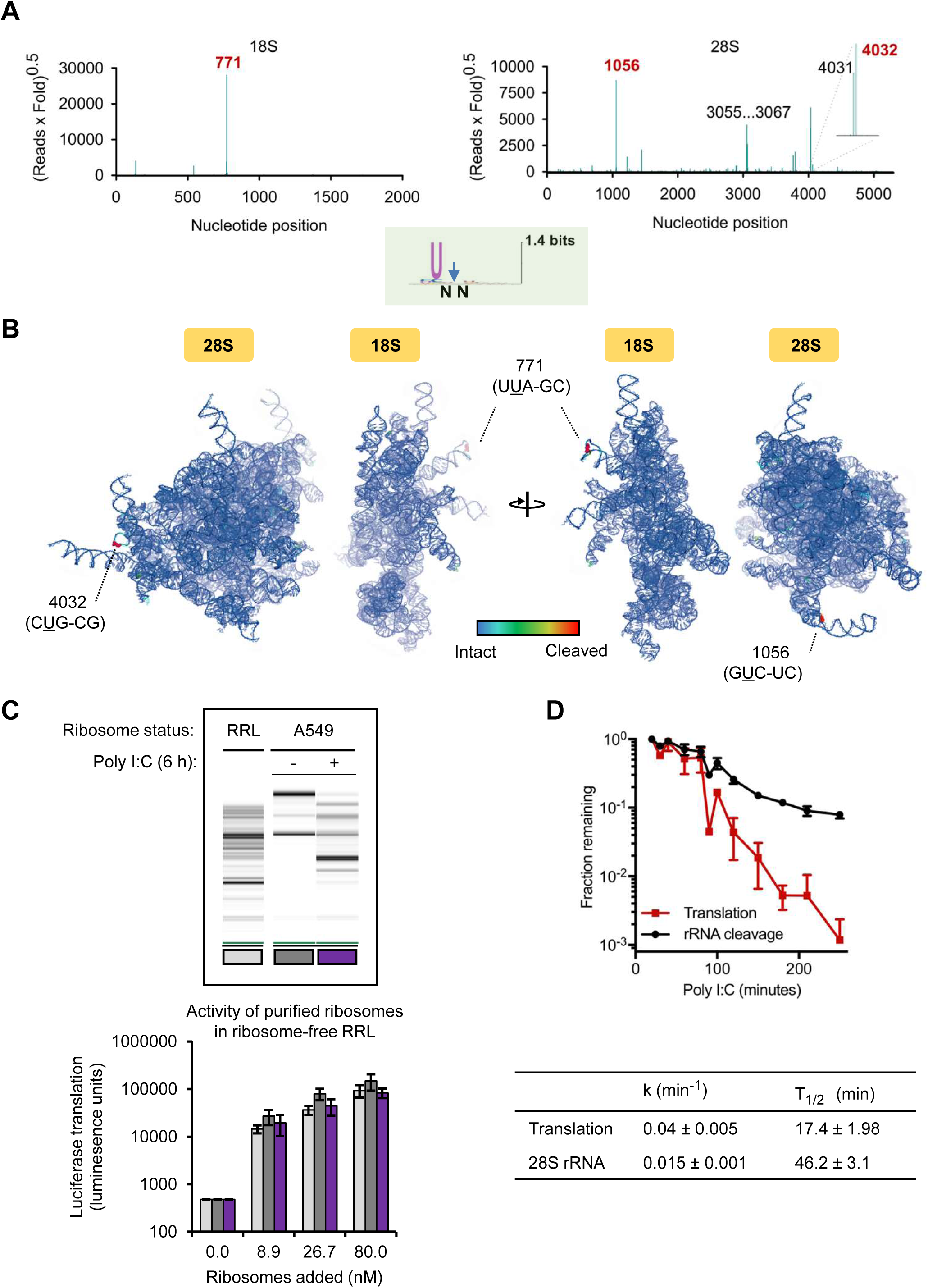
Analysis of ribosomes during 2-5AMD. (A) RNase L cleavage positions in 18S and 28S rRNA found by RtcB RNA-seq. The Y-axis provides a unified metric of read abundance and cleavage induction strength. (B) Mapping the RNase L sites onto 3D structures of 18S and 28S rRNAs. Structures are colored by cleavage strength as defined in (A). (D) Status of rRNA and translation activity of purified ribosomes obtained from rabbit reticulocyte lysate (RRL) or A549 cells. A titration with purified ribosomes was done in the presence of 50 ng capped luciferase mRNA. New translation was measured by luminescence. (D) Comparison of translation loss and rRNA cleavage over the duration of dsRNA response. Fraction of intact rRNA observed by NanoChip was quantified in GelQuant.NET. New translation relative to the untreated condition was measured by ^35^S metabolic labeling and ribopuromycilation (Fig. 1C; S1A). Kinetic parameters of the time profiles are shown below the graph.

Mapping of the identified sites onto the 3D structure of human rRNA shows that except for the nucleotides 4031/4032 in the L1 stalk, 2-5AMD targets surface loops away from vital parts of the ribosome. The location of sensitive sites at distant ribosomal positions suggests that 2-5AMD is not optimized for targeting a defined ribosomal position, as observed with *bona fide* ribosome-inactivating nucleases such as *α*-sarcin (Gluck et al., 1994; Korennykh et al., 2006). The RNase L cleavage sites appear opportunistic rather than intended for ribosomal inhibition.

To test this prediction, we directly assessed the translation activity of the cleaved ribosomes in rabbit reticulocyte lysate (RRL). We depleted this lysate of ribosomes by centrifugation (Fig. S3B) and re-supplied intact ribosomes purified from RRL, A549 cells, or cleaved ribosomes from dsRNA-transfected human cells (Fig. 3C). Using capped luciferase mRNA translation as the readout, we assessed the activity of each ribosome type. In ribosome-depleted RRL, luciferase translation was absent, suggesting that we obtained a suitable assay. Addition of either rabbit ribosomes from nuclease-treated RRL, intact human ribosomes, or human ribosomes with rRNA degraded by 2-5AMD, readily supported translation. We observed the same specific activity for intact and cleaved ribosomes, which we reproduced over a range of ribosomal concentrations to exclude saturation effects (Fig. 3C). Therefore, 2-5AMD does not functionally damage human ribosomes even after a nearly complete degradation of full-length rRNA (Fig. 3C, last lane). In agreement with this observation, the single-exponential decay kinetics for rRNA lags behind the kinetics of translational shutdown (Fig. 3D). Our data and the previously reported disconnect between rRNA cleavage and translation (Donovan et al., 2017) together indicate that the loss of global translation during 2-5AMD involves a process physically distinct from rRNA degradation.

### Spike-in poly-A^+^ RNA-seq Reveals Global Decay of Messenger RNAs during 2-5AMD in Live Cells

In the presence of normal cap-dependent initiation and functional ribosomes, the loss of cell-wide protein synthesis (Fig. 1C, Fig. S1A-B) may arise from cleavage of a non-ribosomal RNA essential for global translation. Cleavage of a tRNA would satisfy this criterion, and tRNA cleavage has been previously demonstrated (Donovan et al., 2017). However, even the most sensitive tRNAs were intact at the time of translational inhibition, suggesting that the only RNA substrates that could account for the translational inhibition are mRNAs. RNase L has been shown to cleave exogenous mRNAs, viral mRNAs and select host mRNAs with a preference for longer RNAs with many AU-rich elements due to the specificity of RNase L for UN^N sites (Al-Ahmadi et al., 2009; Le Roy et al., 2007; Nogimori et al., 2018; Rath et al., 2015). To produce the uniform loss of global translation by mRNA decay, 2-5AMD must act similarly on all housekeeping mRNAs. Indeed, there are no protein bands or protein groups that stand out during the time-dependent progressive loss of translation (Fig. 1C, S1A-B).

To test whether a uniform mRNA decay is taking place, we used qPCR that we designed to detect full-length mRNAs (Fig. 4A). Using this assay we observed a rapid decay of a several abundant basal mRNAs (Fig. 4B). As expected, the decay was absent in RNase L^-/-^ cells (Fig. S4A). All tested mRNAs were cleaved rapidly. The decay traces for all mRNAs leveled before 100% cleavage, suggesting that cells have 2-5AMD-sensitive and 2-5AMD-resistant mRNA pools. The size of the resistant fraction was higher for PRKDC and SON mRNAs (Fig. 4B, dotted lines). By quantifying log-linear regions of the data (Fig. 4B, inset graphs), we determined first-order decay kinetics for each mRNA (Fig. 4C). The housekeeping mRNAs decay considerably faster than rRNA, and on the same time scale as the translational arrest (Fig. 3D vs 4C).

**Figure 4.**
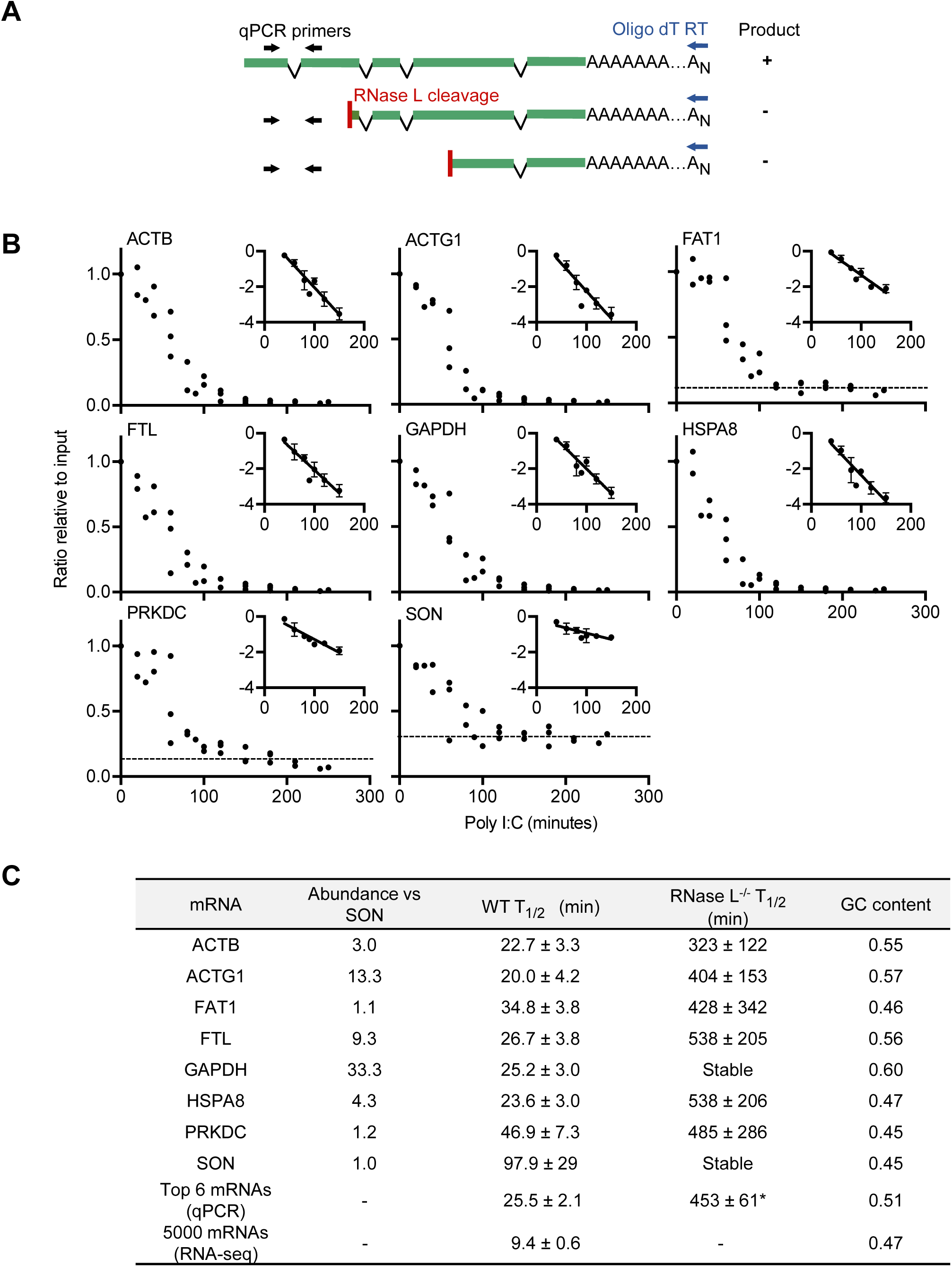
Analysis of decay kinetics for select basal mRNAs during 2-5AMD. (A) Design of qPCR for detection of full-length mRNAs. (B) Decay of select highly expressed housekeeping mRNAs upon poly I:C treatment of A549 cells measured by qPCR. The decay of SON and PKRDC shows distinctly biphasic character indicating the presence of non-cleaved mRNA fraction (2-5AMD-resistant pool) in the cells. The inset graphs show log-linear parts of the decay profiles used to measure the first-order decay kinetic parameters. Dotted lines mark approximate levels of 2-5AMD-resistant mRNA fractions. (C) Decay half-life (T_1/2_) obtained for each mRNA from the qPCR in (B). The last line shows T_1/2_ for 5,000 mRNAs calculated based on RNA-seq (Fig S4B). (*) Excluding GAPDH due to its stability.

Using poly-A^+^ RNA-seq we extended our analysis to the transcriptome. Widely used RNA-seq normalization and differential expression analysis techniques presume that levels of most mRNAs remain unchanged. This assumption would be violated if 2-5AMD inhibited translation by global mRNA decay. To correctly quantify mRNA levels during the course of decay, we supplemented our RNA samples with internal standards (*Drosophila Melanogaster* RNA spike-ins). 2-5AMD profiling revealed a time-dependent loss of almost all cellular mRNAs (Fig. 5A; Tables S3). The RNA-seq data agreed well with our qPCR analysis (Fig. 5B). Reads for all decaying mRNAs disappeared across the entire transcript length, suggesting that once RNase L endonucleolytically cleaves a transcript, the resulting mRNA fragments are rapidly cleared. The decay kinetics determined from the top 5000 most abundant transcripts matches the time scale of translational inhibition (Fig. S4B; 4C). Together, our qPCR and RNA-seq data link translation arrest by 2-5AMD to mRNA decay, which can explain the accumulation of empty 80S monosomes in cells.

**Figure 5.**
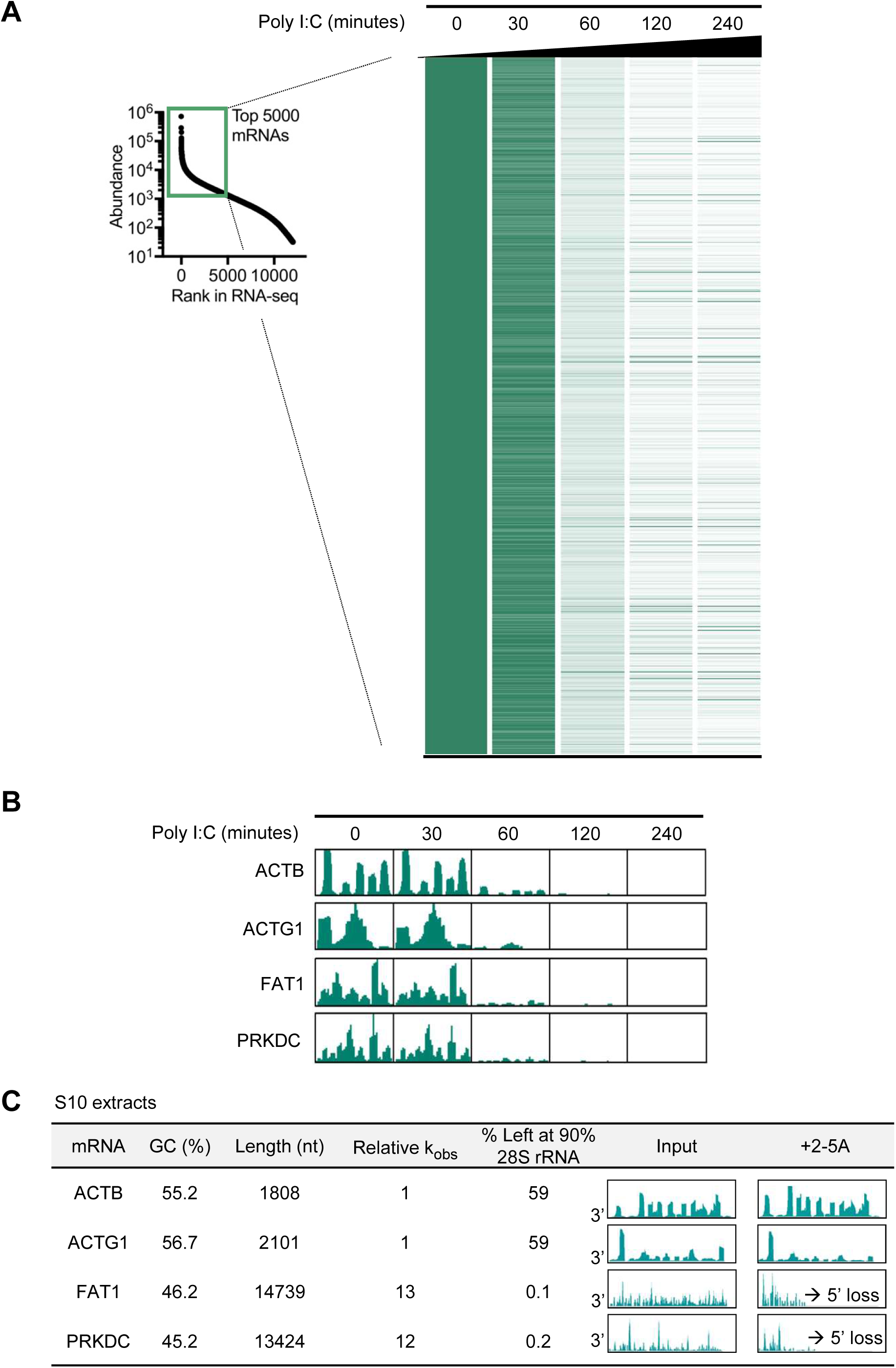
RNA-seq profiling of mRNA decay by RNase L in live cells and in cytosolic extracts. (A) Time-dependent decay of the most abundant 5,000 mRNAs measured by spike-in RNA-seq. To obtain mRNAs levels, total reads for each sample were normalized using *D. melanogaster* RNA spike in as an internal standard. Transcripts are ordered from the highest to the lowest expression level in the untreated sample. (B) RNA-seq profiles for select individual mRNAs from (A). (C) Cleavage profiles and kinetic parameters for the mRNAs in (B) obtained in cell-free experiments (Rath et al., 2015).

Of note, evaluation of RNase L activity in cytosolic cell extracts showed that mRNA decay depends on both, mRNA length and AU content (Rath et al., 2015) (Fig. S5). In S10 cytosolic extracts treated with 2-5A, at the time point when ∼60% ACTB mRNA still remains, only 0.1% of FAT1 mRNA and 0.2% of PRKDC mRNAs survive (Fig. 5C). The high sensitivity of FAT1 and PRKDC transcripts in the S10 extract is in line with their lower GC content and greater length leading to more net UN^N sites per mRNA. In contrast to these findings, 2-5AMD in live cells shows no dependence on GC content and leads to decay of PRKDC, FAT1, ACTB and most other mRNAs with comparable, within several-fold, kinetics (Fig. 5A-B; Discussion). Therefore, cellular mRNA decay agrees with the fast timing of translational inhibition and the loss of global protein synthesis.

### Decay and Synthesis Kinetics Protect IFN mRNAs from 2-5AMD

Spike-in RNA-seq indicates that the 2-5AMD-sensitive RNAs that decay (Fig. 6A, 88-89% of poly-A^+^ RNA) account for more than 99.7% of protein synthesis (Fig. 1C, S1A). These data suggest that RNase L eliminates the most actively translating mRNAs. As much as 11-12% of the poly-A^+^ RNA is resistant (Fig. 6A), indicating that resistant mRNAs must be shielded from RNase L, perhaps by being in the nucleus, in phase-separated complexes or in stress granules. The pool of RNase L-resistant poly-A^+^ transcripts is enriched with non-coding RNAs (ncRNAs; Fig. 6A), suggesting that translation could render mRNAs more sensitive to RNase L compared to ncRNAs. The potential dependence on translation appears to be in line with the recently proposed model for translation-dependent RNase L cleavage based on analysis of decay of exogenous RNA (Nogimori et al., 2018).

**Figure 6.**
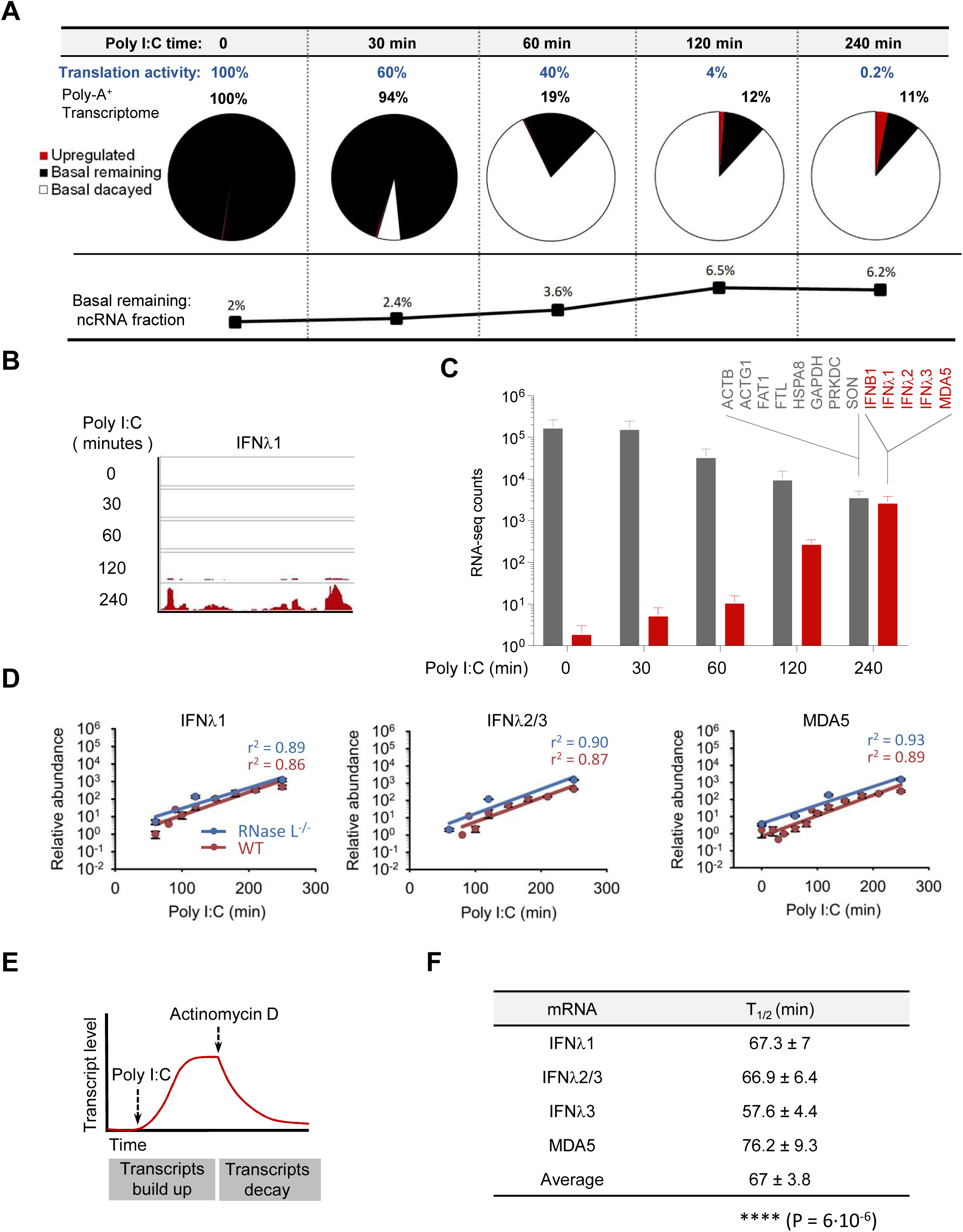
Poly-A^+^ transcriptome composition and dynamics during 2-5AMD. (A) Time evolution of the poly-A+ transcriptome obtained from spike-in RNA-seq. Induced transcripts (red) were upregulated by ≥ 5-fold at four hours of poly I:C treatment. (B) Illustration of read increase for a gene that is induced. (C) Levels of select basal and defense mRNAs measured by spike-in RNAs-seq. (D) Induction of innate immune mRNAs in response to poly I:C with and without 2-5AMD. (E) Experimental outline for measurement of innate immune mRNA decay. Innate immune mRNAs are allowed to accumulate followed by transcriptional inhibition with actinomycin D and decay profiling. (F) Kinetics parameters of defense mRNAs measured as in Fig. 4B. P-value compares T_1/2_ for defense vs basal mRNAs.

As basal mRNAs decay, defense mRNAs encoding IFNs and ISGs are upregulated (Fig. 6A). The mRNAs are initially absent, but by four hours IFN/ISG mRNAs account for 25% of the poly-A^+^ pool. In log-linear coordinates, the loss of basal mRNAs and the increase in innate immune transcripts obeys a linear law (Fig. 6C, S7A). The induction of defense transcripts bypasses RNase L (Fig. 6D; S6A; S7A), as previously reported (Alisha Chitrakar et al., 2018). To test whether defense mRNAs could survive in the presence of RNase L, we measured their decay kinetics during 2-5AMD in the presence of actinomycin D treatment, which was used to stop new transcription (Fig. 6E). RNase L-dependent decay of the innate immune mRNAs was readily detected (Fig. 6F; S6B), indicating that defense mRNAs can be cleaved by RNase L. However, the decay half-lives (T_1/2_) for defense mRNAs were ∼2-3-fold longer compared to those of basal mRNAs (Fig. 6F).

The log-linear decay of basal mRNAs indicates that 2-5AMD degrades these transcripts with single-exponential kinetics. In contrast, the log-linear increase of IFN/ISG mRNAs does not match single-exponential accumulation or steady influx laws, and indicates induction with a positive feedback (Fig. S7B; S7A-D; Methods). In accord with our data, the presence of positive feedback has been previously described for IFN signaling (Michalska et al., 2018). It is important to note that positive feedback in the IFN response involves not only RNase L-sensitive mediators (IFN/ISG mRNAs), but also RNase L-resistant activators (accumulation of IFN proteins and phosphorylation of the transcription factor STAT) (Fig. 7A). We developed a mathematical description of this positive feedback model to examine whether it predicts survival of defense mRNAs in the presence of RNase L. To define experimental parameters, we used the key observations that (1) basal mRNAs are downregulated by ∼100-fold after 4 hours, (2) 2-5AMD has a small effect on the dynamics of defense mRNAs, and (3) measured half-lives of defense mRNAs are longer than half-lives of basal mRNAs (Fig. S7A; Fig. 6F).

**Figure 7.**
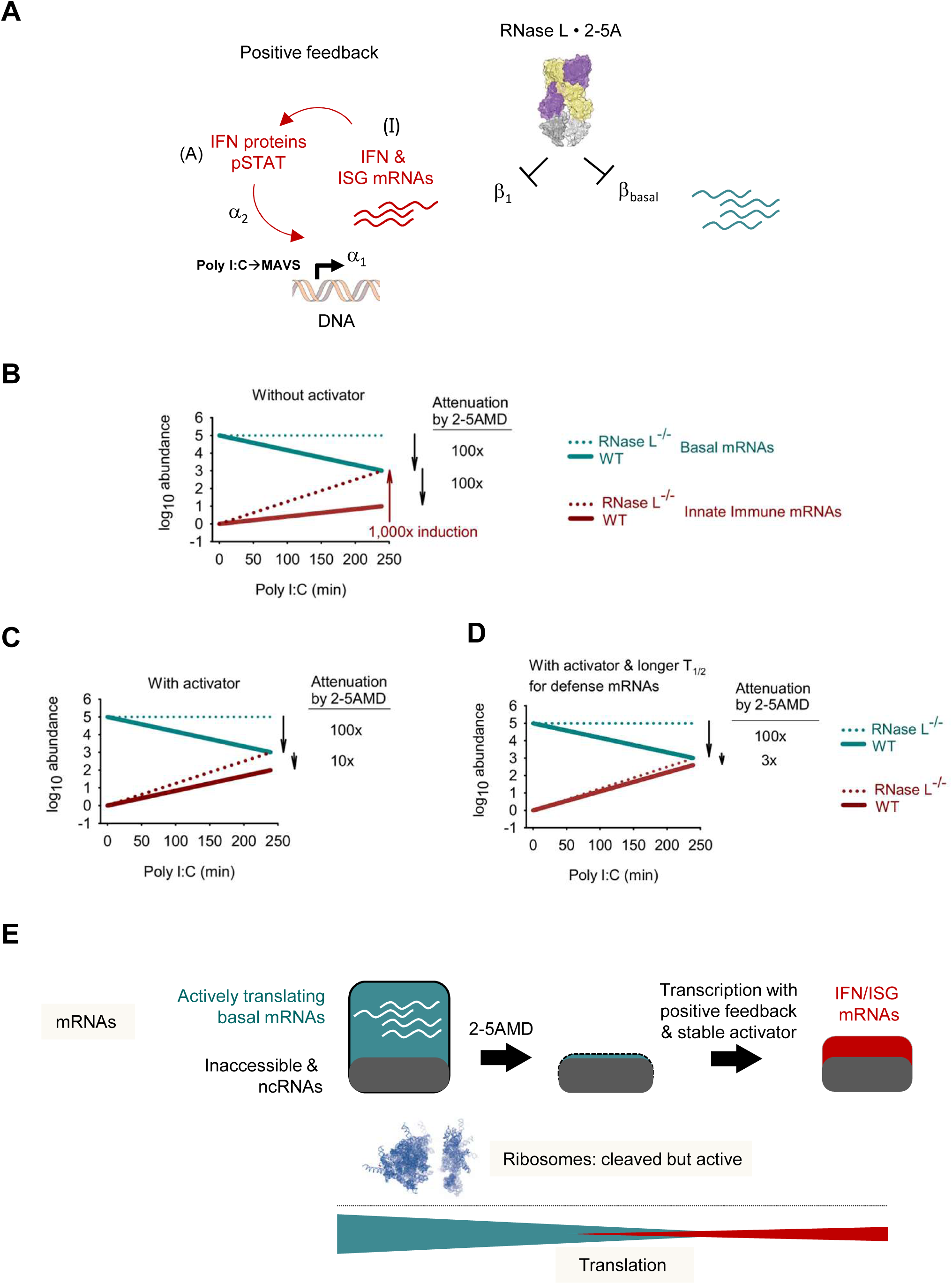
Proposed model for coordination between transcription and decay in 2-5AMD. (A) A kinetic scheme for 2-5AMD and the IFN response with positive feedback. Parameters I (IFN mRNAs), A (2-5AMD-resistant stable activator within positive feedback), α and β are described in “Decay and transcriptional dynamics analysis section” (Methods). (B) Modeled dynamics of basal and defense mRNAs when all transcripts have the same sensitivity to 2-5AMD. Defense mRNAs are induced with positive feedback. (C) Model in (B) modified to include a stable activator of positive feedback. (D) Model in (B) modified to include both, a stable activator and stability of defense mRNAs. The modeling analysis in panels A-C is described in Methods. (D) The proposed model for mRNA decay and IFN transcriptional response leading to RNase L-mediated reorganization global translation.

A simple model with positive feedback and similar 2-5AMD sensitivity of all mRNAs predicts equal attenuation by RNase L of basal and transcriptionally induced mRNAs (Fig. 7B). Therefore, positive feedback *per se* is not protecting defense mRNAs from 2-5AMD. In contrast, protection is observed once a stable activator is included in the positive feedback loop (Fig. 7C). The protection arises from the inherent resistance of proteins (IFNs, phosphorylated STAT) to RNase L. IFN proteins steadily accumulate and promote increasing transcriptional activity even in the presence of 2-5AMD. The evasion from 2-5AMD is further enhanced once 2.6-fold greater mRNA stability is incorporated in the model (Fig. 7D). Our modeling results show that longer half-lives of defense mRNAs, as well as, independently, a positive feedback loop built around a stable activator can result in evasion of defense mRNAs from global decay.

## Discussion

Our work suggests that in mammalian cells decay of abundant mRNAs can be efficiently coupled with a kinetically matched transcriptional response to steer translation toward stress proteins. Since the discovery of translational inhibition by 2-5AMD in 1977 (Hovanessian et al., 1977), various hypotheses have been proposed to explain how this mechanism contributes to the IFN response and how mammals may use 2-5AMD to cope with stress conditions associated with dsRNA. The hypothesized functions ranged from antiviral defense due to viral RNA decay (Cooper et al., 2014a; Han et al., 2004; Nilsen and Baglioni, 1979) to IFN amplification by cleaved self-RNAs (Malathi et al., 2007) or destruction of host RNAs as a mechanism to kill infected cells during the late pro-apoptotic stages of dsRNA sensing (Chakrabarti et al., 2011; Zhou et al., 1997). The observations that 2-5AMD can begin early and precede the IFN response, and that IFNs bypass 2-5AMD (Alisha Chitrakar et al., 2018; Donovan et al., 2017) revealed unanticipated translational reprogramming in this pathway, leading to the present investigation. Our model, based on analysis of 2-5AMD in human A549 cells, is summarized in Figure 7F. We show that the loss of protein synthesis can be explained by depletion of thousands of host mRNAs. The same mechanistic conclusion is made in a manuscript submitted back to back with our work (Burke et al., and Roy Parker [ref will be added]; See not added on submission). This group too attributed the action of RNase L to global loss of mRNAs. We show that approximately 90% of the total mRNA pool (by mass) is sensitive to 2-5AMD and accounts for nearly the entire translational activity detected by both metabolic and ribopuromycilation assays. The remaining 11-12% of the poly-A^+^ transcriptome is neither accessible to RNase L nor supporting translation, suggesting that RNase L may have a mechanism to preferentially target translationally-active endogenous mRNAs.

RNase L cleaves most basal mRNAs with similar rate constants and does not exhibit a preference for longer and AU-rich mRNAs. The uniform mRNA decay is a central feature of 2-5AMD, which is responsible for arrest of all housekeeping proteins rather than just proteins encoded by long and AU-rich mRNAs. We propose that this uniformity indicates that in live cells mRNAs are cleaved in Briggs-Haldane kinetic regime (Methods). Under Briggs-Haldane conditions, RNase L will cleave mRNAs independently of the binding (K_m_) and catalytic constants (k_cat_) of individual mRNAs, thereby acquiring a mechanism for uniform decay of the transcriptome. A notable feature of Briggs-Haldane regime is that once RNase L encounters an mRNA, it makes a cut before dissociating, i.e. every RNase L-mRNA binding event is productive. In cell extracts, RNase L is sensitive to length and AU content in mRNAs, indicating cleavage under Michaelis-Menten regime. Although it remains to be explained precisely how live cells achieve Briggs-Haldane regime, our observations could be explained by ribosome-assisted RNase L access to mRNAs. This model agrees with the recently proposed mechanism for Dom34 rescue of RNase L-ribosome complex, which postulates a translation-dependent mRNA cleavage mechanism (Nogimori et al., 2018). If ribosome-RNase L recognition (rather than mRNA-RNase L recognition) determines kinetics mRNA decay by RNase L, 2-5AMD would target all actively translating mRNAs. Further, RNase L could dwell on translating ribosomes, which would ensure efficient cleavage of mRNAs and achieve Briggs-Haldane conditions.

The decay rate constants are similar, within several-fold, for basal mRNAs and for mRNAs encoding IFNs (Fig. 4C; 6F). We show that once this several-fold stability advantage is coupled with the positive feedback of the IFN response, defense mRNAs become desensitized to RNase L. A minimal model with experimentally determined kinetic parameters can account for decay of basal mRNAs and explain how IFNs and ISGs can accumulate to nearly the same levels in WT and RNase L^-/-^ cells (Fig. 7A-B). The experimental observation that 2-5AMD can clear the cytoplasm from unneeded host mRNAs without compromising the innate immune system suggest that, by analogy, 2-5AMD could also eliminate viral mRNAs while allowing the IFN response to develop. The antiviral activity of RNase L poses a serious obstacle for some viruses, which evolved dedicated viral proteins to disarm 2-5AMD usually by degrading 2-5A or binding to RNase L (Drappier et al., 2018; Gusho et al., 2014). Our data raise the possibility that some viruses may alternatively rely on replication kinetics as a strategy of escape from the innate immune action of 2-5AMD, without using inhibitory viral proteins. Understanding whether viruses take advantage of such a kinetic mechanism will be important for understanding and improvement of antiviral treatments.

Reprogramming protein synthesis is a central strategy by which mammalian cells achieve energy conservation and adapt to stressful environments. To our knowledge, the mechanism of prioritizing stress protein translation by 2-5AMD is different from mechanisms of previously described mammalian pathways. Whereas mammalian protein synthesis is usually regulated by interference with translation initiation factors (Iwasaki et al., 2016; Marques-Ramos et al., 2017; Taniuchi et al., 2016), 2-5AMD acts directly on messenger RNAs. If global mRNA decay is matched with a complementary transcriptional response, the transcriptional induction can outpace the decay. In the described example, activation of this mechanism orchestrates cell commitment to the IFN response.

### Database entries

RNA-seq data have been submitted to GEO database under accession ID GSE123034.

## Acknowledgments

The authors thank Prof. Kevan Shokat for the gift of INK128 and helpful discussions, Dr. Paul Copeland (Rutgers University) for the gift of internal ribosome entry site (IRES)-containing dual luciferase plasmid, staff at the Proteomics and Mass Spectrometry Core facility, and staff at the Genomics Core Facility at Princeton University. We are grateful to Prof. Zemer Gitai and to Dr. Sophia Li for help with polysome analysis, to Prof. Bonnie Bassler, Prof. Martin Jonikas, and Prof. Sabine Petry for sharing instrumental resources and we thank members of the Korennykh lab for great help throughout the project.

## Author contributions

S.R. and E.P. conducted the core experimental work. J.D. and K.D. conducted select experiments. N.S.W. and Y.M. conducted theoretical analysis of decay and transcription kinetics. S.R., E.P. and A.K. wrote the manuscript. A.K. supervised the work.

## A note added on submission

Our work was submitted to BioRxiv back to back with a related and independent study by Burke et al., and R. Parker [ref will be added here]. Our work employs a considerably different set of experimental approaches, but is synergistic with the co-submitted paper. Our manuscripts converge on a similar model of mRNA regulation by 2-5AMD.

## Supplementary Information

**Figure S1.**
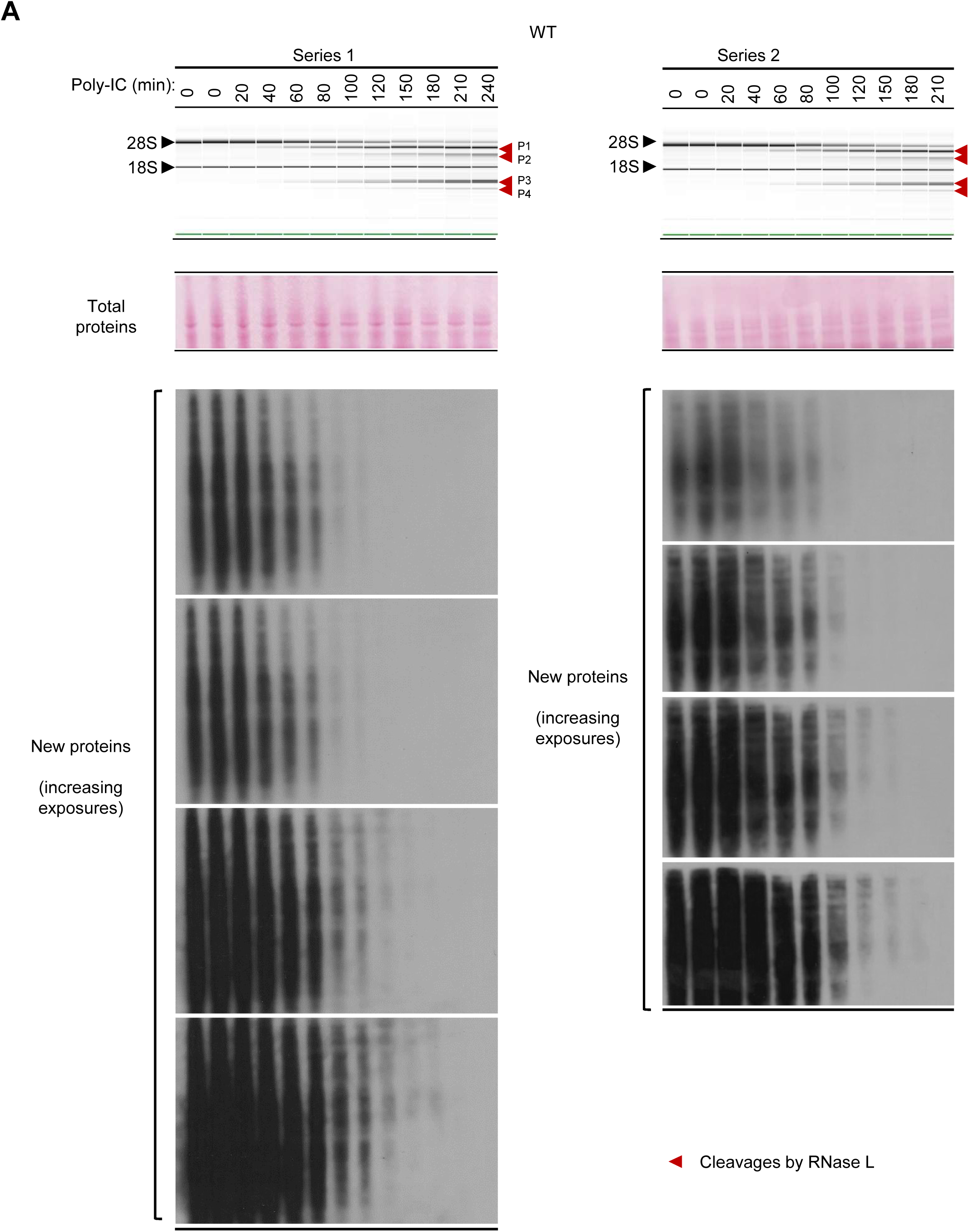
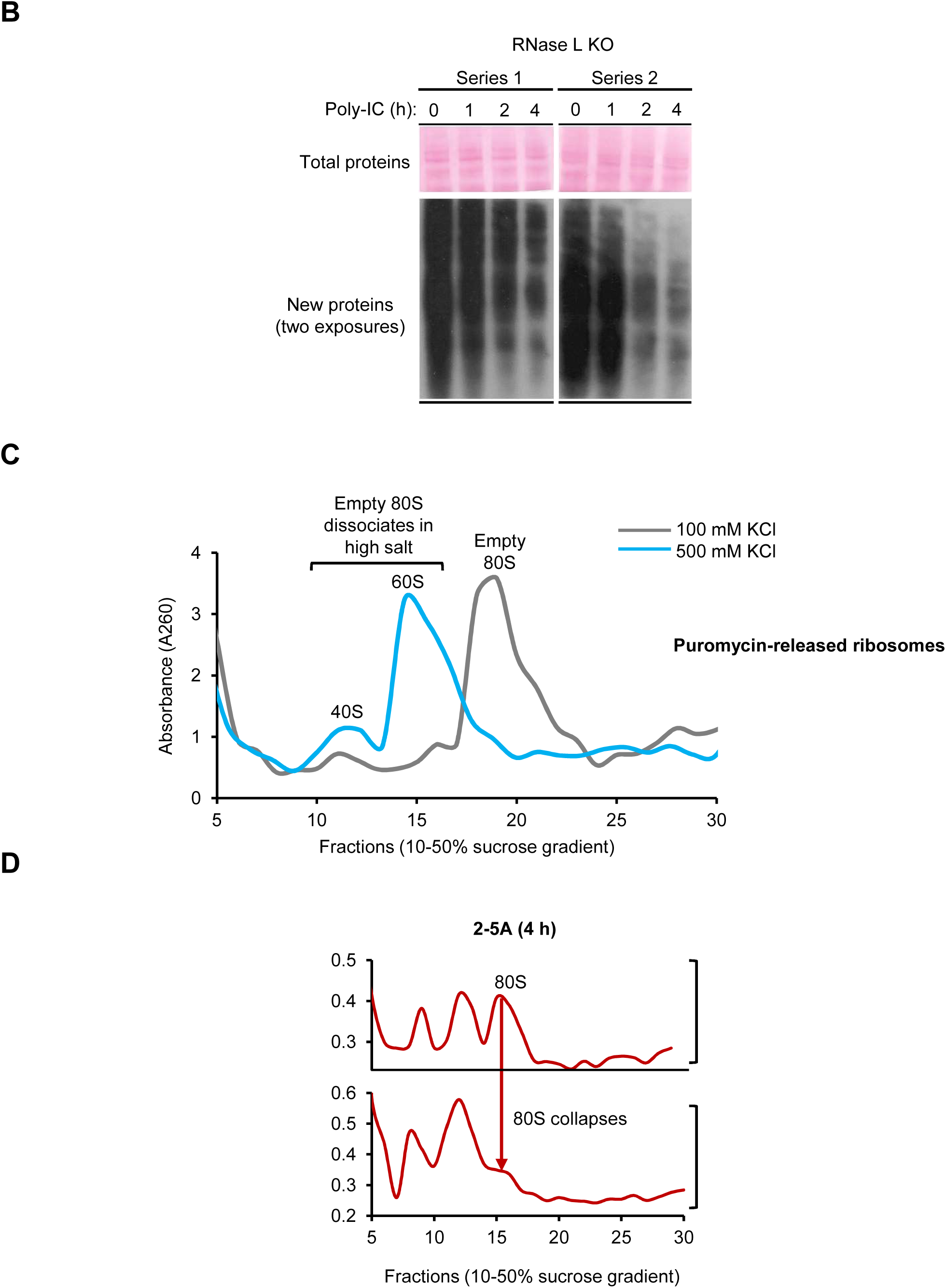
Translation regulation by 2-5AMD. (A) Dynamics of rRNA cleavage by 2-5AMD (monitored by NanoChip) vs nascent protein synthesis monitored by puromycin labeling of nascent peptides. Western blots against puromycin show ribopuromycilated peptides arising from active translation (Methods). Multiple exposures are shown to more fully capture the dynamic range. Ponceau stain provides loading control. (B) Translation in RNase L^-/-^ cells complementary to data in (A). (C) Puromycin-released naïve ribosomes analyzed on a sucrose gradient containing 100 mM vs 500 mM KCl. Puromycin-released ribosomes are not bound to mRNA and migrate as empty 80S in 100 mM KCl. These empty ribosomes dissociate into 40S and 60S species in 500 mM KCl. (D) CHX-stalled ribosomes from naïve and 2-5A-treated cells analyzed by sucrose gradient at 100 mM vs 500 mM KCl.

**Figure S2.**
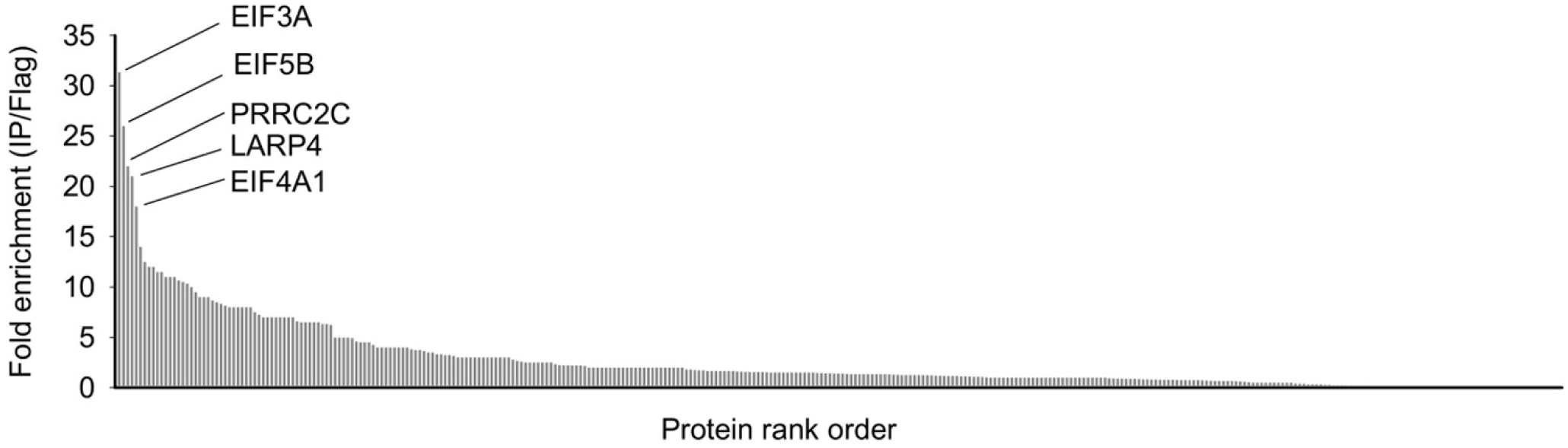
Key proteins identified by mass spectrometry of eIF4E pulldown. The mass-spectrometry data sets are provided in Table S1.

**Figure S3.**
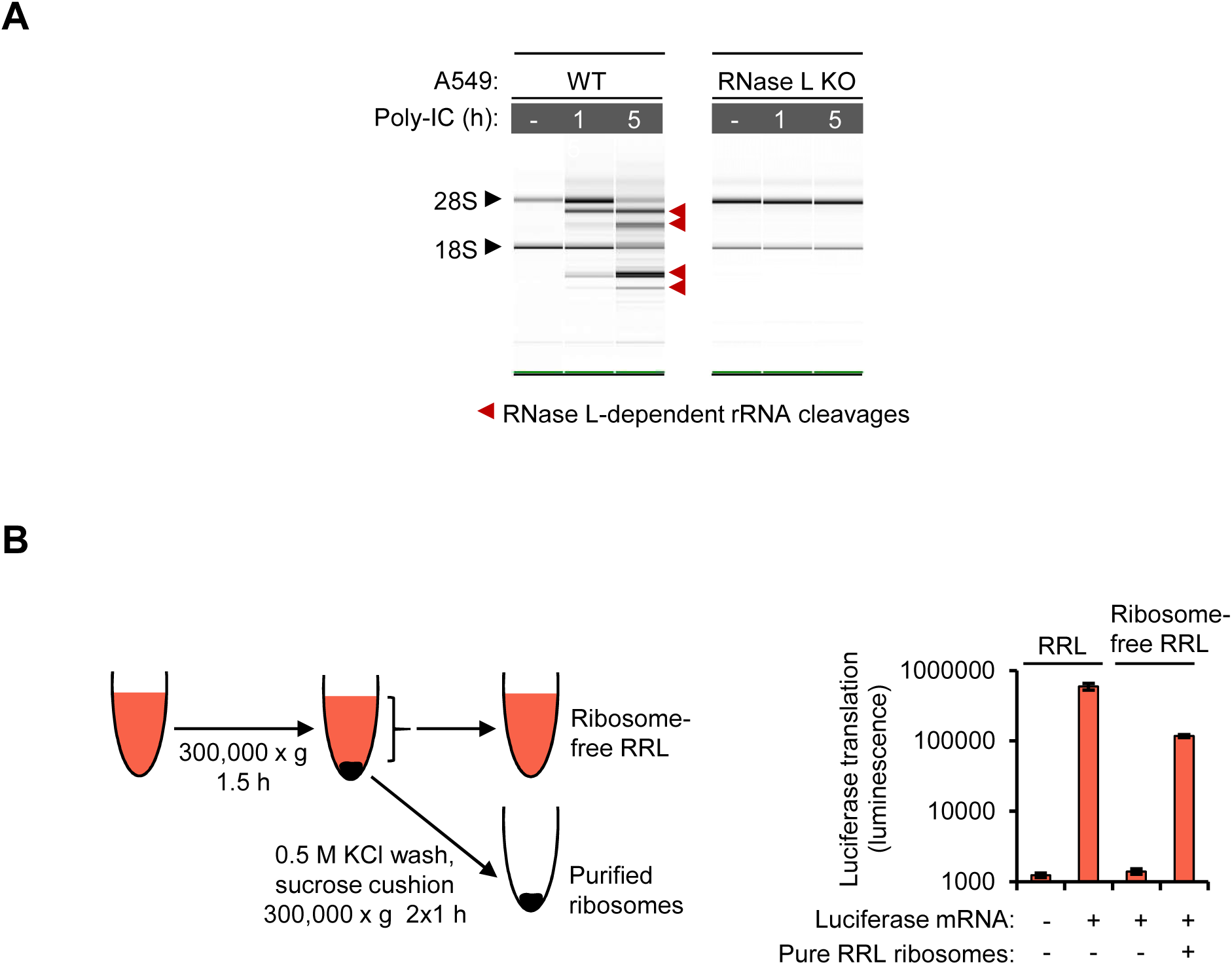
Ribosome cleavage and translation analysis assays. (A) NanoChip profiles of rRNA samples used for mapping RNase L sites by RtcB-seq. (B) Schematic outline for preparation of ribosome-free RRL and its validation using translation activity assay. Translation of 50 ng capped luciferase mRNA was conducted in the presence of 80 nM RRL ribosomes. The translation activity was measured by luminescence (Methods).

**Figure S4.**
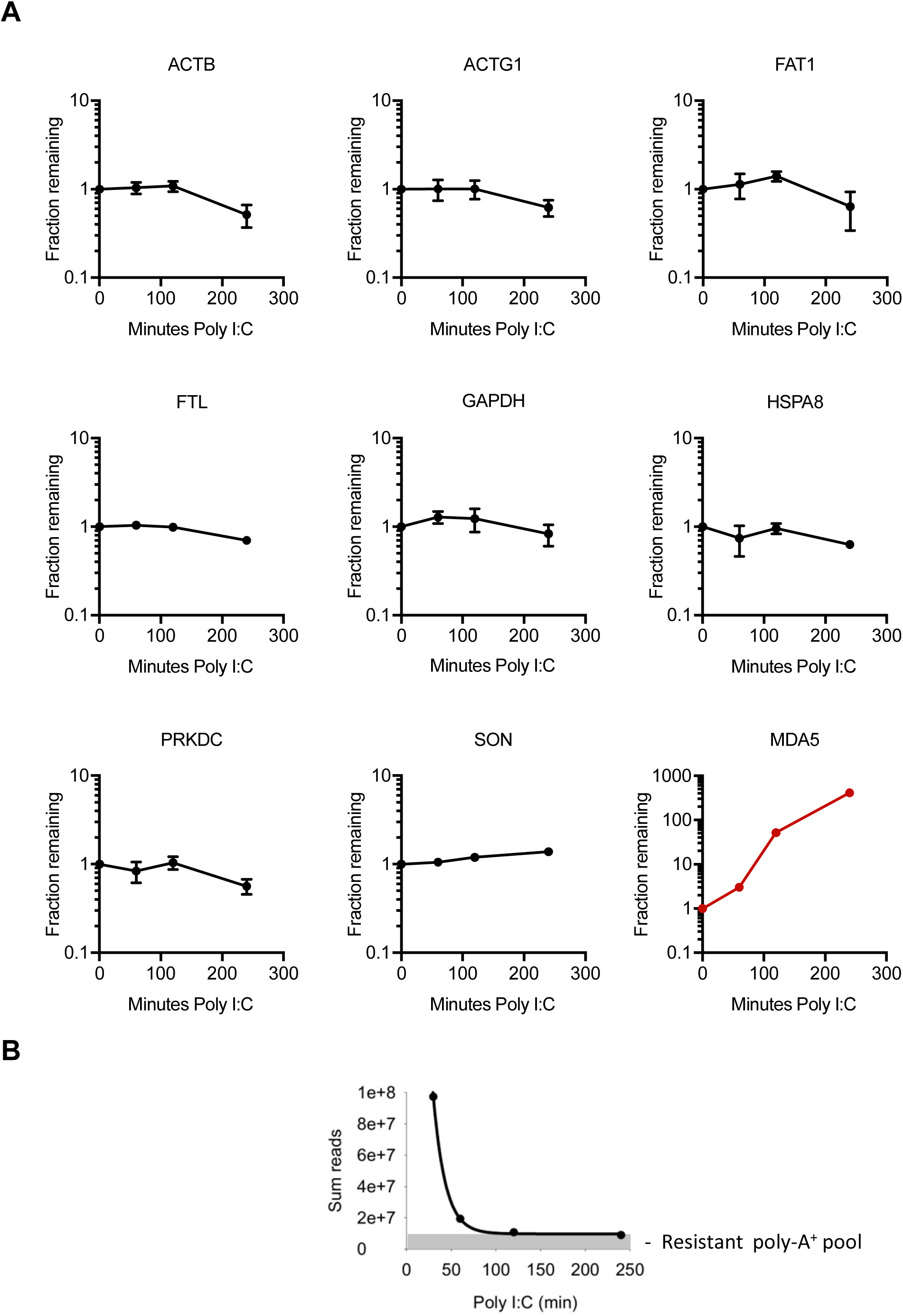
Messenger RNA Dynamics in cells treated with poly I:C. (A) The levels of basal mRNAs determined by qPCR using primers designed as in Fig. 4A. Traces for housekeeping mRNAs are colored black. The trace for an IFN-stimulated gen, MDA5, is colored red. Error bars show S.E. from three qPCR replicates. (B) Single-exponential decay of top 5000 mRNAs during 2-5AMD. Experimental values represent summed read counts determined by poly-A^+^ RNA-seq, normalized using *Drosophila Melanogaster* RNA spike-ins.

**Figure S5.**
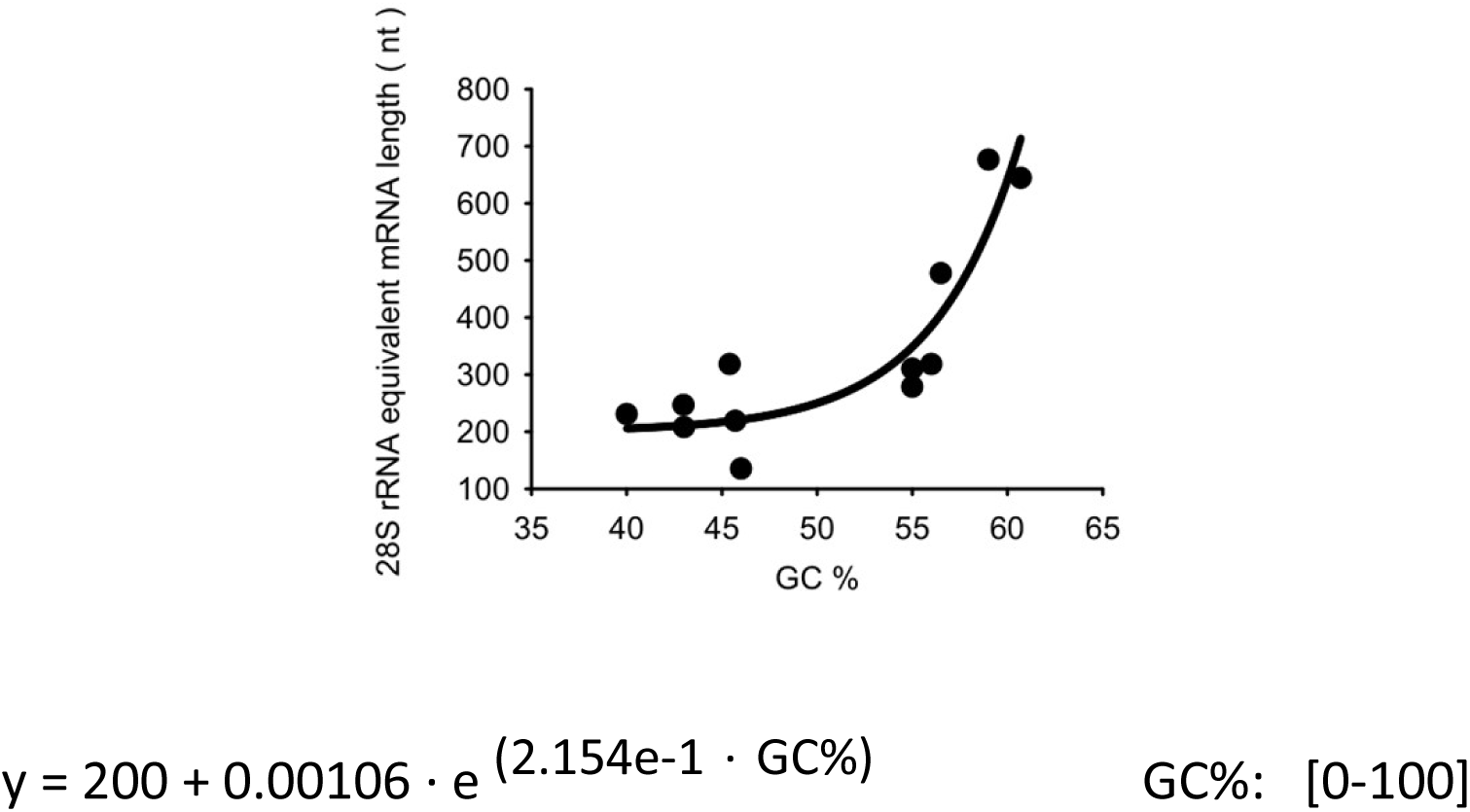
GC composition and length sensitivity of mRNAs to RNase L in cell-free systems. In cytosolic extracts, length-normalized sensitivity of different messenger RNAs depends primarily on their GC content, as illustrated by the graph. The GC can be used to calculate “28S rRNA-equivalent mRNA length” (REML), defined as mRNA length that decays with the same rate as 28S rRNA. Application of REML for calculation of mRNA decay in cell extracts is described in Methods. The data were obtained from RNA-seq GEO dataset GSE75530 (Rath et al., 2015).

**Figure S6.**
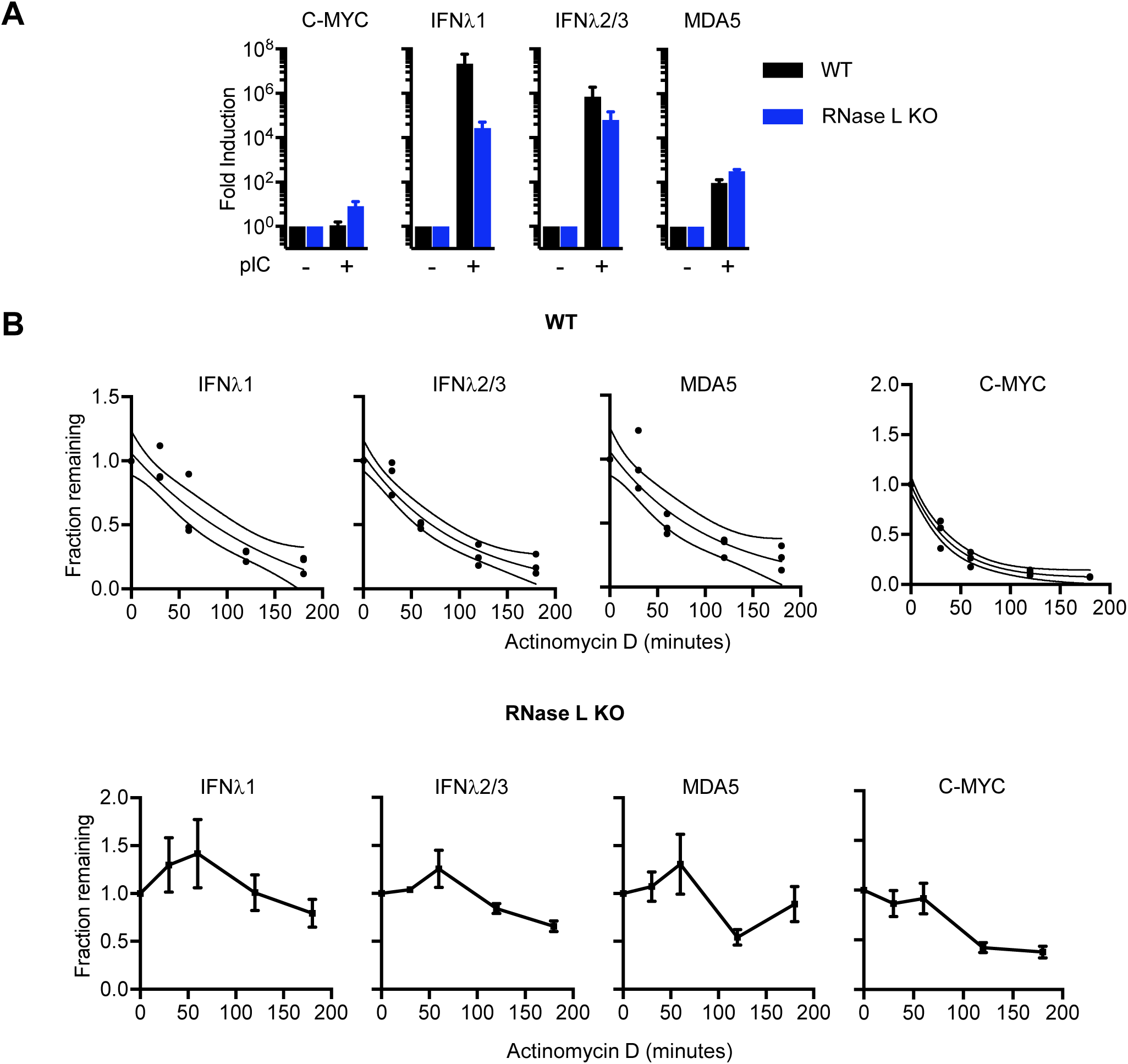
Decay kinetics of innate immune mRNAs. (A) Induction of innate immune mRNAs by poly I:C measured by qPCR. (B) Decay analysis of innate immune mRNAs in WT and RNase L-/-A549 cells. Following activation of the IFN response and 2-5AMD, cells were treated with actinomycin D to arrest new transcription. Decay of innate immune mRNAs was subsequently monitored by qPCR. A short-lived non-immune mRNA encoding c-MYC is shown for reference.

**Figure S7.**
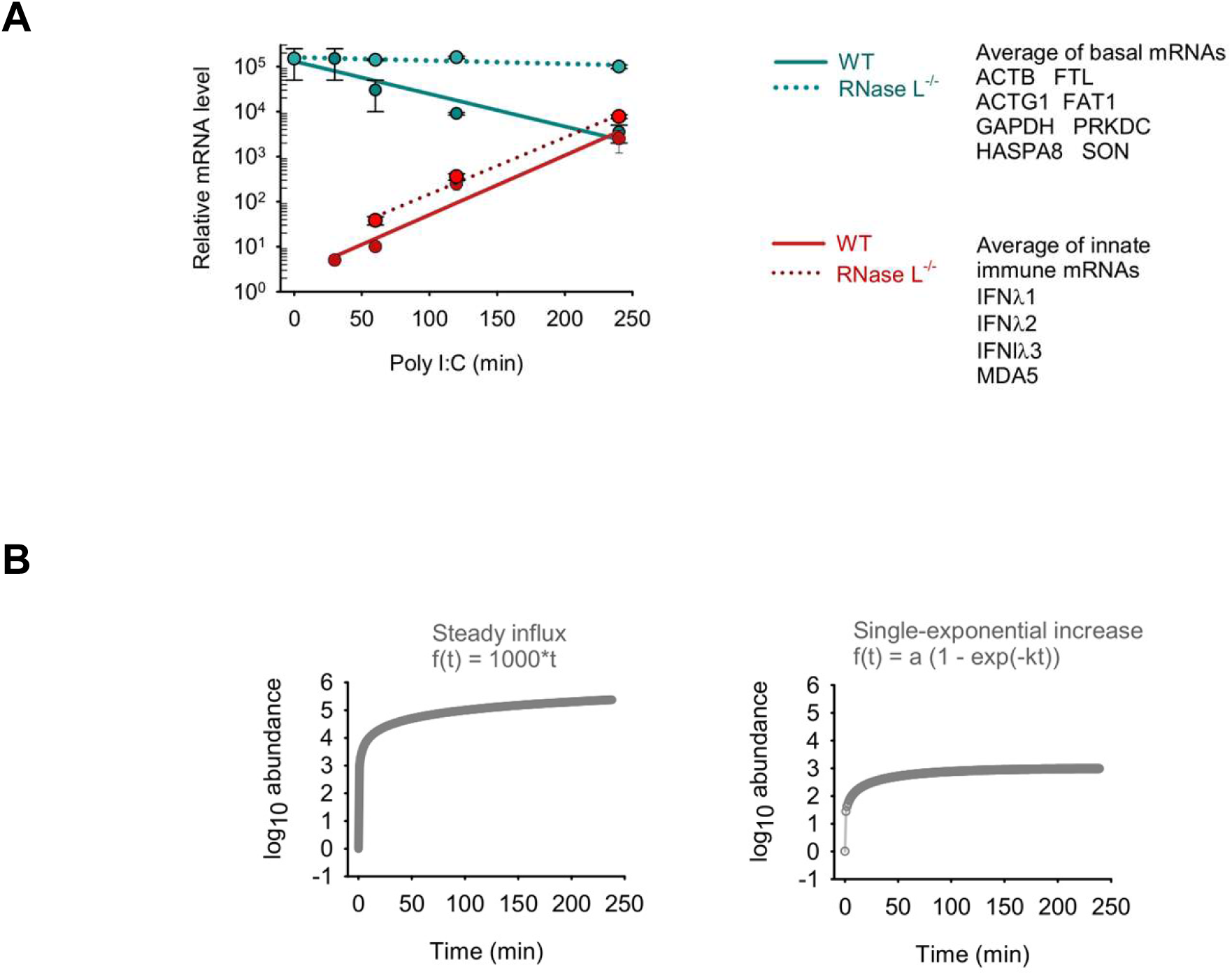
Modeling mRNA dynamics in dsRNA response. (A) Integrated data from figures 4B, 6C and 6D. (B) Alternative models of IFN/ISG induction. Steady influx model (left): IFN is produced at a steady rate. First-order kinetics model (right): IFNs increase according to single-exponential law.

## Methods

### Cell culture

Human cells were grown in RPMI with 10% FBS. For poly I:C transfections, 1 µg/mL poly I:C was transfected using Lipofectamine 2000 Reagent (Thermo Fisher Scientific) for the indicated durations. For experiments aimed at measuring mRNA decay rates, RNA polymerase II transcription was blocked by adding 1 µg/ml actinomycin D directly to the cell culture medium for the indicated durations. To measure decay rates of poly I:C-induced transcripts, WT and RNase L^-/-^ cells were treated with 1 µg/ml poly I:C for 4 hours, followed by actinomycin D treatment.

### Nascent translation analysis by ^35^S and ribopuromycilation

To conduct ^35^S metabolic labeling, cells were incubated in methionine-free RPMI (Gibco) containing 11 µCi EasyTag EXRESS35S Protein Labeling mix (Perkin Elmer) for 15 minutes at 37 °C. Cells were directly lysed in NuPage LDS sample buffer. Lysates were boiled at 95 °C for 10 minutes, then separated on 10% BisTris PAGE gels (Invitrogen). Gels were stained with coomassie for total protein visualization, then analyzed by phosphorimaging (Typhoon FLA 7000, GE). For ribopuromycilation assay, cells were treated with 10 µg/ml puromycin (Invitrogen) in culture medium for 5 minutes at 37 °C. Cells were lysed and separated by PAGE as above. For western blotting proteins were transferred to PVDF membranes (Life Technologies) and stained with Ponceau Red to visualize total proteins. The membrane was washed and blocked in 5% non-fat dry milk in TBST. Membranes were probed with 1:1000 dilution of mouse anti-puromycin antibody (EMD Millipore), followed by 1:10,000 dilution of horseradish peroxidase-conjugated anti-mouse secondary antibody (Jackson ImmunoResearch).

### RtcB RNA-seq

RtcB RNA-seq was conducted as described previously (Donovan et al., 2017), but without short RNAs purification step. Briefly, 1 µg RNeasy-purified RNA was ligated to 10 μM adaptor (Table 1, oligo 1). Ligation reactions were performed using 10 μM RtcB, 20 mM HEPES pH 7.5, 110 mM NaCl, 2 mM MnCl_2_, 100 μM GTP, 40U RiboLock RNase inhibitor (Fermentas), 4 mM DTT, and 0.05% Triton X-100 for 1 hour at 37 °C. Reactions were quenched using 3 mM EDTA. Owing to its short length, free adapter was removed by purifying the ligated RNA with the RNeasy kit. On-column DNase treatment was omitted so that ligated adapter remained intact. Oligonucleotides were reverse-transcribed using MultiScribe reverse transcriptase (RT) and 2 pmol of RT primer complimentary to the ligation adaptor (Table 1, oligo 2). RNA, RT primer, and dNTPs were incubated for 3 min at 65°C and snap-cooled on ice. MgCl_2_ (3 mM f/c) was added to ensure efficient Mg^2+^-dependent RT reaction.

**Table 1.**
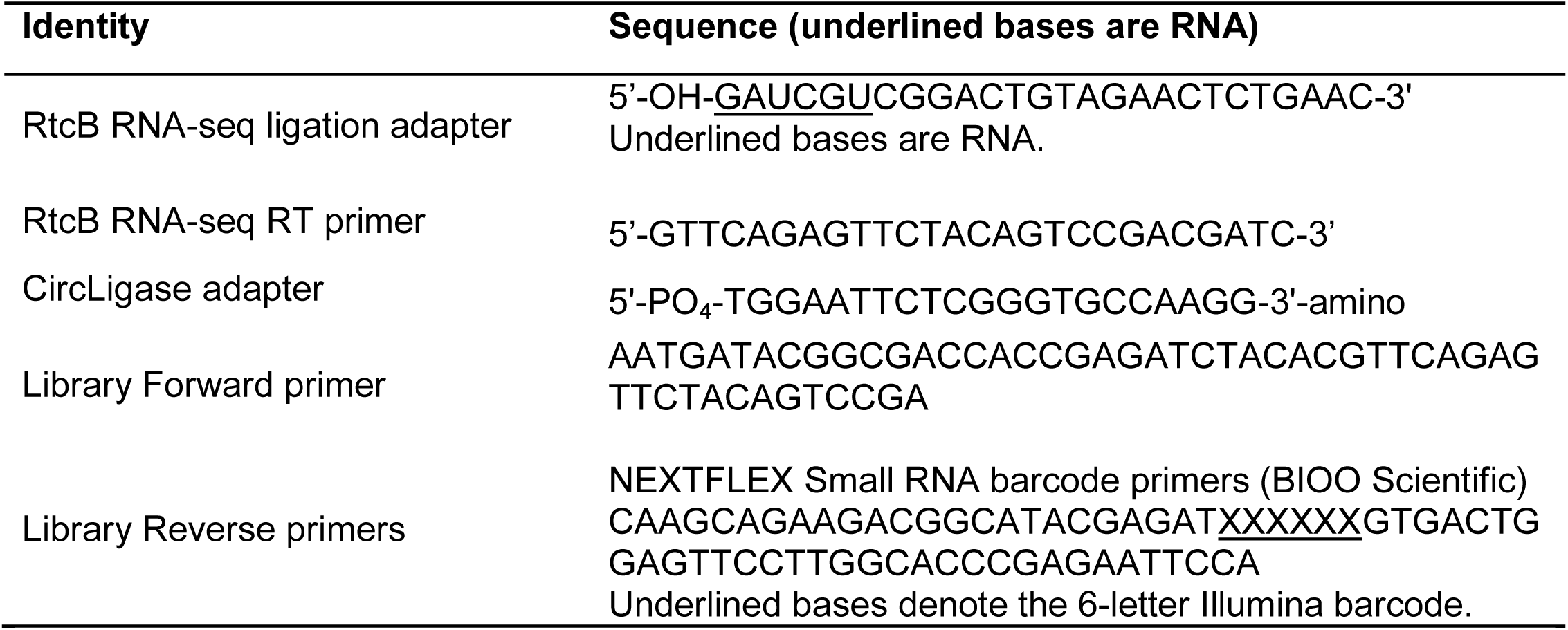
Oligonucleotides used for RtcB RNA-seq library preparation.

A 2x mastermix containing RT buffer, RT (Applied Biosystems), and 40U Ribolock was added to snap-cooled samples to a final volume of 20 μL. Reactions were incubated at 25°C for 10 min, 37°C for 1.5 h, and 95°C for 5 min. cDNA reaction was brought up to 200 μl with water and extracted using 1 volume of 25:24:1 Phenol:Chloroform:Isoamyl Alcohol saturated with 10 mM TRIS (pH 8.0) and 1 mM EDTA (Sigma Aldrich). The aqueous phase was precipitated using 20 μg glycogen as a carrier, 2/3 5M ammonium acetate (vol/vol) and 3 volumes of 100% ethanol. Mixture was incubated at –80 °C for 30 minutes, followed by a 17,000 x g spin at 4 °C, for 30 minutes. Pellets of cDNA were washed with 75% ethanol (vol/vol) and resuspended in 25 μl DI water. Reactions contained 30% of the cDNA from the previous step, 1 μM adaptor (Table 1, oligo 3), 1 U/μl CircLigase (Epicentre), and buffer contents as per manufacturers guidelines. CircLigase reactions were incubated for 1 h at 65°C and quenched by adding EDTA to a final concentration of 8 mM. 1/3 of the quenched CircLigase reaction was PCR amplified using Phusion DNA polymerase (NEB) and primers 4-5 in Table 1. Libraries were analyzed by a BioAnalyzer high sensitivity DNA 1000 chip (Agilent). Equimolar amounts were pooled and gel purified from native page. RtcB RNA-seq was performed on an Illumina HiSeq 2500 and processed as described previously (Donovan et al., 2017).

### Poly-A^+^ RNA-seq

RNA purified by RNeasy kit (1 μg) was used for poly-A^+^ enrichment with oligo-dT beads. Pulldown was followed by standard fragmentation, adapter ligation and PCR amplification for sequencing on the Illumina HiSeq 2000 platform. Sequencing reads were mapped to the human genome hg19 using TopHat 2 (Kim et al., 2013), set to map stranded reads with default parameters. Reads mapping to exons of each gene were counted using HTseq-count in union mode (Anders et al., 2015). RNA-seq data were visualized using the Integrative Genomics Viewer (Robinson et al., 2011).

### Polysome sedimentation analysis

Cells in 10 cm dishes were flash frozen in liquid nitrogen, scraped in cold PBS with 100 µg/ml CHX and pelleted at 500 x g for 5 minutes at 4°C. The cell pellet was lysed in 5 mM HEPES, 1.5 mM KCl, 2.5 mM MgCl_2_, 100 µg/ml CHX, 1x Protease inhibitor cocktail, 100 U/mL RNase inhibitor (NEB), 0.5 % Triton X-100, and 0.5 % Na-Deoxycholate. The lysate was vortexed, rotated end-over-end for 7 minutes at 4°C and centrifuged at 10,000 x g for 10 minutes at 4 °C. Clarified lysate was layered over a 12 ml 10-50 % sucrose gradient made by GradientMaster (BioComp). The 10 % and 50 % sucrose solutions were made with 20 mM HEPES, 100 mM KCl, 5 mM MgCl_2_, 100 μg/mL CHX, 1x Protease inhibitor cocktail, and 100 U/ml RNase inhibitor. In experiments designed to distinguish mRNA bound 80S vs empty 80S complexes, both lysis and sucrose gradient buffers were adjusted to a final concentration of either 100 mM or 500 mM KCl. To create empty 80S as a control, WT cells were pre-treated with 50 μg/mL puromycin for 20 min and the sucrose gradient buffer also contained 50 μg/ml puromycin in place of CHX. The lysate was spun through the gradient in an SW41Ti rotor in an Optima XE-100 Ultracentrifuge (Beckman Coulter) at 200,000 x g for two hours at 4 °C. BioComp Gradient Fractionator was used to fractionate the gradients and the 254 nm absorbance was read by an EM-1 ultraviolet monitor (BioRad).

### qPCR

RNA was purified using the RNeasy kit (Qiagen). RNA quality was assessed using BioAnalyzer NanoChip (Agilent) and extent of rRNA cleavage was quantified using GelQuant.NET (Biochem Lab Solutions, http://biochemlabsolutions.com/). Within each experiment, equal amounts (ng) of RNA were converted to cDNA using oligo-dT_18_ as primer and the High Capacity Reverse Transcriptase kit (Applied Biosystems). qPCR was performed using Power SYBR Green PCR Master Mix (Life Technologies).

### eIF4E immunoprecipitation

Magnetic protein A beads were incubated with 2 µg anti-FLAG (Sigma) or anti-eIF2α (Santa Cruz) antibodies in IP buffer (10 mM HEPES (pH 7.5), 150 mM NaCl, 0.1% Nonidet P-40 (NP-40), 1x complete protease inhibitor cocktail, RNase inhibitor) at 4°C for two hours. Excess unbound antibody was removed by washing the beads twice in IP buffer for 5 minutes. Cells transfected with or without 2-5A for three hours were lysed in 10 mM HEPES (pH 7.5), 10 mM NaCl, 2 mM EDTA, 0.5% Triton X-100, 1x complete protease inhibitor cocktail and RNase inhibitor for 7 minutes while rotating, at 4 °C. Lysates were clarified at by spinning at 10,000 x g for 10 minutes and incubated with antibody-bound beads for two hours at 4 °C. After two hours, beads were subject to 5 x 2 min washes with IP buffer. RNA and protein from inputs, supernatants (unbound) and IPs were analyzed using qPCR, western blot and mass spectrometry, respectively.

### Mass spectrometry

Gel bands were digested using 1.5 µg Trypsin (Promega). Samples were dried completely in a SpeedVac and resuspended with 21 µl of 0.1% formic acid (pH 3). Next, 5 µl was injected per run using an Easy-nLC 1200 UPLC system. Samples were loaded directly onto a 45 cm long 75 µm inner diameter nano capillary column packed with 1.9 µm C18-AQ (Dr. Maisch, Germany) mated to metal emitter in-line with an Orbitrap Fusion Lumos (Thermo Scientific, USA). The mass spectrometer was operated in data dependent mode with the 120,000 resolution MS1 scan (AGC 4e5, Max IT 50ms, 400-1500 m/z) in the Orbitrap followed by up to 20 MS/MS scans with CID fragmentation in the ion trap. Dynamic exclusion list was invoked to exclude previously sequenced peptides for 60s if sequenced within the last 30s and maximum cycle time of 3s was used. Peptides were isolated for fragmentation using the quadrupole (1.6 Da) window. Ion-trap was operated in Rapid mode with AGC 2e3, maximum IT of 300 ms and minimum of 5000 ions.

Raw files were searched using Byonic (Bern et al., 2012), MS-Amanda (Dorfer et al., 2014) and Sequest HT algorithms (Eng et al., 1994) within the Proteome Discoverer 2.2 suite (Thermo Scientific, USA). 10 ppm MS1 and 0.4 Da MS2 mass tolerances were specified. Carbamidomethylation of cysteine was used as fixed modification, oxidation of methionine, acetylation of protein N-termini, conversion of glutamine to pyro-glutamate and deamidation of asparagine were specified as dynamic modifications. Trypsin digestion with maximum of two missed cleavages were allowed. Files searched against the Uniprot *Homo Sapiens* database downloaded on February 23, 2017 and supplemented with common contaminants. Scaffold (version 4.8.4, Proteome Software Inc., Portland, OR) was used to validate MS/MS based peptide and protein identifications. Peptide identifications were accepted if they could be established at greater than 90.0% probability by the Scaffold Local FDR algorithm. Protein identifications were accepted if they could be established at greater than 99% probability and contained at least 2 identified peptides. Protein probabilities were assigned by the Protein Prophet algorithm (Nesvizhskii et al., 2003). Proteins that contained similar peptides and could not be differentiated based on MS/MS analysis alone were grouped to satisfy the principles of parsimony.

### *In vitro* transcription

An internal ribosome entry site (IRES)-containing dual luciferase plasmid was a gift from Dr. Paul Copeland (Rutgers University). Monocistronic firefly luciferase was obtained by PCR amplification of the coding region from the dual luciferase construct and cloning into BamHI/NotI digested pcDNA3.1. Plasmids were linearized with AgeI (firefly luciferase) or BamHI (dual luciferase) and purified by phenol extraction and ethanol precipitation. Capped mRNAs were transcribed using reagents from the MEGA ShortScript Kit, except for nucleoside triphosphates, and 12 mM anti-reverse cap analog (NEB). NTPs were added using a custom 10X mixture containing 75 mM each of ATP, UTP, and CTP, and 15 mM GTP. Transcription was carried out for two hours at 37 °C followed by addition of Turbo DNase I and incubation for 20 minutes at 37 °C. Messenger RNAs were phenol extracted and purified on P30 micro spin columns (Bio-Rad).

### Ribosome-depleted Rabbit Reticulocyte Lysate (RRL)

Ribosome-free RRL was prepared essentially as described (Gupta et al., 2013). Briefly, nuclease treated RRL (Promega) was centrifuged 2 x 1 hour at 300,000 x g, 4 °C with care to not disturb the ribosome pellet when removing the supernatant. The pellet from the first centrifugation was saved for purifying RRL ribosomes from the salt-wash step.

### Ribosome purification

Frozen A549 cell pellets (∼200 µL) were resuspended in 500 µL of 20 mM HEPES-KOH (pH 7.5), 100 mM KCl, 5 mM MgCl_2_, 4 mM DTT, 0.2% NP-40, 1x phosphatase inhibitors 2/3 (Sigma), 2x complete protease inhibitor (Roche), and 0.4 U/ml RNase inhibitor (NEB), as described previously (Lorsch and Herschlag, 1999). Resuspended cells were rotated for 15 minutes at 4°C, followed by centrifugation for 20 minutes at 16,000 x g, 4°C. Obtained supernatants were centrifuged for 30 min at 21,000 x g, 4°C. The resulting supernatants were centrifuged for 80 minutes at 300,000 x g and 4 °C to yield a crude ribosomal pellet. Pellets were washed with 5 mM HEPES-KOH (pH 7.5), 50 mM KCl, 1.5 mM MgCl_2_, 4 mM DTT and resuspended in 100 µL of the fresh wash buffer. KCl was adjusted to 0.5 M and ribosomes were incubated for additional 30 minutes on ice, followed by centrifugation for 5 minutes at 10,000 x g and at 4 °C to remove debris. Salt-washed ribosomes (130 µl) were layered onto a 100 µl 0.5 M sucrose cushion containing 20 mM HEPES-KOH (pH 7.5), 100 mM KCl, 2 mM MgCl_2_, and 4 mM DTT. Tubes were centrifuged for 90 minutes at 300,000 x g, at 4°C and the obtained pellets were rinsed with 3 x 50 µL of the ribosome storage buffer (20 mM HEPES KOH pH 7.5, 50 mM KCl, 2 mM MgCl_2_, 4 mM DTT, 10% glycerol) and then dissolved. Debris was removed by centrifugation as above. A small quantity of purified ribosomes (1 μl) was processed with Trizol for rRNA extraction. Remaining ribosomes were quantified, aliquoted, and flash-frozen in LN2. Ribosomes were quantified using 5×10^7^ M^−1^cm^−1^ as the molar extinction coefficient.

### Cell-free translation analysis

Cell-free translation experiments were conducted using nuclease-treated rabbit reticulocyte lysate (Promega). Reactions were 12.5 µl and contained 8 µl RRL (ribosome-depleted or not, as indicated), 0.25 µl 40 U/ml RNase inhibitor, 0.25 µl 1 mM amino acids, 50 ng capped firefly luciferase mRNA or 300 ng dual luciferase mRNA (3 µl combined volume of mRNA and H_2_O), and 1 µl ribosome storage buffer or ribosomes to achieve the indicated final ribosome concentrations. For firefly luciferase mRNA, reactions were incubated for 30 minutes (firefly luciferase mRNA) at 30 °C and then quenched by adding 50 µl 20 mM HEPES pH 7.5, 100 mM NaCl and 1 mM MgCl_2_. The terminated reactions were transferred to a 96-well plate and supplemented with 10 µl of 6X luciferin mix (20 mM HEPES pH 7.5, 100 mM NaCl, 36 mM MgCl_2_, 2.4 mM D-luciferin, 18 mM ATP). Luminescence was measured for 10 seconds using a Tristar2 Multi-Mode Plate Reader (Berthold Technologies).

### Decay and transcriptional dynamics analysis

The main observation is that while basal mRNAs decay, the rate of increase of IFNs is only slightly reduced in the presence of RNase L (Fig. S7A). The simplest model to describe the dynamics of IFN mRNAs is that, in the absence of RNase L, they increase exponentially due to direct positive feedback according to the rate equation:

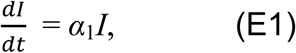

with the solution:

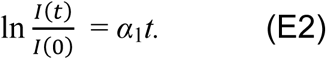

With RNase L, IFN mRNA loss due to decay can be accounted for by adding a decay rate constant *β*_1_:

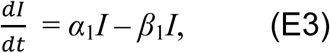

With solution:

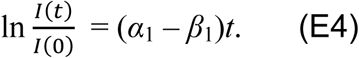

However, the observed decrease of the rate of accumulation of IFN mRNAs due to the presence of RNase L is much smaller than this model would predict (Fig. S7A).

In order to explain the observation that the rate of increase of IFNs is only slightly reduced in the presence of RNase L, we generalize the above model by assuming that the IFN positive feedback loop is mediated by a stable activator (e.g. the IFN protein and phosphorylation of the transcription factor STAT) which do not get cleaved by RNase L. They provide a gradually accumulating activator, increasing stimulating IFN mRNA transcription with time. Denoting by *I*(*t*) and *A*(*t*) the concentrations of IFN mRNAs and the IFN proteins, respectively, the rate equations that describe the system become:

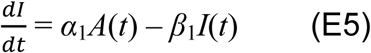

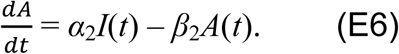

RNase L does not degrade *A*, therefore we set *β*_2_ = 0. The solution of these equations is given by 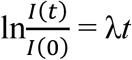 where λ is given by

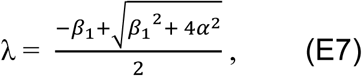

where we only consider the relevant increasing solution and define 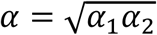. In the absence of decay of the IFN mRNAs (*β*_1_ = 0), one obtains λ = *α*, i.e. IFN induction depends only on IFN mRNA induction and activator induction but not on RNase L. If IFN mRNA decay is present and *β*_1_ << *α* (which is the case, see Fig. S7A), then λ ≈ *α* – *β*_1_/2. Under these conditions, the effect of RNase L on IFN accumulation will be a contribution to decay at just 1/2 the potency of the bare rate of RNase L-mediated decay of IFN mRNA. For decay of basal mRNA, *β*_basal_ = 0.007 (in natural logarithm scale and with units ∼1/time). The innate immune mRNA decay ∼2.6-fold slower on average, i.e. *β*_1_ ∼ 0.003. The effect of a stable activator will attenuate this value 2-fold to give λ ≈ *α* – 0.0015. Considering that decay-free IFN accumulation occurs with α ∼ 0.013 (Fig. S7A), subtraction of 0.0015 will have a negligible effect, which explains why RNase L does not strongly inhibit IFN production. A graphical representation of these results is provided in Figures 7B-D.

For more general parameter values, note that if we define *γ* = *β*_1_ / *α*, we can write 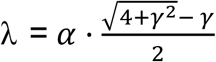, which is always positive. Based on this relationship, even if the decay exponent *β*_1_ is very large (i.e. 2-5AMD is very strong), as long as a stable activator is present in the positive feedback loop there will be exponential growth of IFNs due to the activator gradually accumulating and leading to faster IFN mRNA synthesis.

Lastly, we note that if the activator does decay (*β*_2_ ≠ 0), the growth exponent is given by the expression

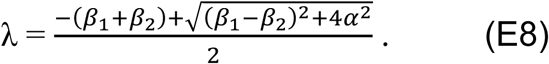

In particular, when *β*_1_ = *β*_2_ (IFN mRNA and the activator both decay at the same rate), λ = *α* - *β*_1_, and the kinetics then reproduces the case of IFN mRNA decay at full potency (E4).

### Application of ribosome-equivalent messenger RNA length (REML) for calculation of mRNA decay in cell-free systems

Using the RNA-seq data (GEO ID GSE75530) we determined that fraction of intact mRNA left in a cell-free system in the course of 2-5AMD can be calculated from mRNA length L, GC content, and 28S rRNA cleavage observed by NanoChip as follows:

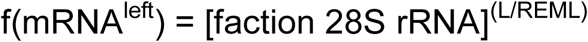

In the case of ACTB (mRNA length L = 1808 nt and GC content of 55.2%), REML = 350 nt (Fig. S5). Using the expression for f(mRNA^left^), it is determined that under 2-5AMD conditions that degrade 10% of 28S rRNA, 58% of ATCB mRNA will be remaining: (0.9)^(1808/350)^ = 0.58. Under conditions of 50% 28S rRNA cleavage, (0.5)^(1808/350)^ = 0.028 (2.8%) of ACTB mRNA will remain. The stability of mRNAs will vary with mRNA length and GC composition. For a transcript with 40% GC (REML = 200 nt) and length 10,000 bases, under conditions of 10% of 28S rRNA degradation only (0.9)^(10000/200)^ = 0.005 (0.5%) of the uncleaved mRNA will be remaining. Therefore under conditions of 10% 28S rRNA cleavage, the latter mRNA will appear to be ∼ 100-fold more sensitive to RNase L than mRNA of ACTB.

### Error analysis

For decay rates, data points from three biological replicates were plotted together. Statistical significance (P) was from Welch two-tailed unpaired t-test (James McCaffrey implementation, Microsoft, https://msdn.microsoft.com/en-us/magazine/mt620016.aspx). *P≤0.05, **P≤0.01, ***P≤0.001, *P≤0.0001, NS: non-significant. Error bars represent S.E.

### Briggs-Haldane kinetics application to 2-5AMD

Michael-Menten kinetics requires rapid enzyme-substrate binding equilibrium E+S. In contrast, in Briggs-Haldane regime the enzyme-substrate binding equilibrium is not achieved because product formation from the ES complex is faster than ES complex dissociation to give free E+S. Under Michaelis-Menten conditions, enzyme preference for two different substrates (specificity) is defined as the ratio: S1/S2, whether S1 is k_cat_^1^/K_m_^1^ for substrate 1 and S2 is k_cat_^2^/K_m_^2^ for substrate 2. Catalytic activity (k_cat_) and binding (K_m_) both determine the relative reaction rates of the two substrates. In Briggs-Haldane regime, the rate constant for the product formation from ES (k_cat_) is much larger than the rate constant for the ES complex dissociation (k_off_), such that k_cat_/K_m_ = k_cat_/(k_off_+k_cat_/k_on_) ∼ k_on_. The specificity ratio S1/S2 is simplified to k_on_^1^/k_on_^2^. S1/S2 no longer depends on the binding affinity (K_m_) or the catalytic activity (k_cat_) and depends only on the association rate constant (k_on_). For similar substrates such as different mRNAs k_on_ will depend primarily on the hydrodynamic radius and should be similar for most mRNAs. If association is mediated by the ribosome, k_on_ for different mRNAs could be determined by ribosome-mRNA or ribosome-RNase L association.

## Tables

**Table 2.**
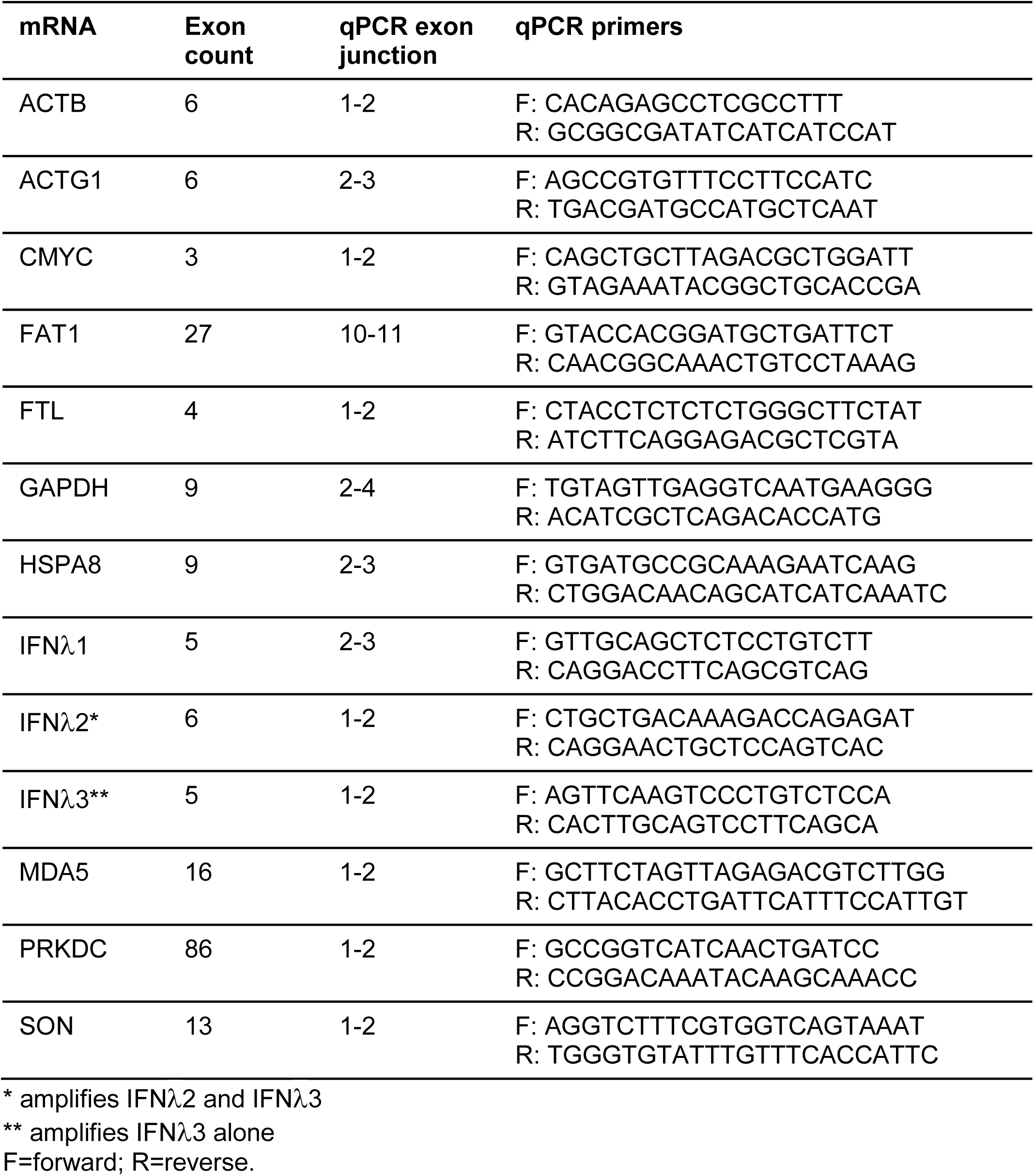
qPCR Primers used in this work.

